# A systems-based map of human brain cell-type enriched genes and malignancy-associated endothelial changes

**DOI:** 10.1101/528414

**Authors:** Philip Dusart, Björn M Hallström, Thomas Renne, Jacob Odeberg, Mathias Uhlén, Lynn M Butler

## Abstract

Changes in the endothelium of the cerebral vasculature can contribute to inflammatory, thrombotic and malignant disorders. The importance of defining cell type-specific genes and how they are modified in disease is increasingly recognised. Here, we developed a bioinformatics-based approach to identify normal brain cell-enriched genes, using bulk RNAseq data from 238 normal human cortex samples from 2 independent cohorts. We compared endothelial cell-enriched gene profiles with astrocyte, oligodendrocyte and neuron profiles. Global modifications to the endothelium in malignant disease were characterised, using RNAseq data from 516 human lower grade gliomas and 401 human glioblastoma multiforme samples. Lower grade glioma appeared to be an ‘endothelial intermediate’ between normal brain and glioblastoma multiforme. We identify the most highly glioblastoma multiforme-specific endothelial cell biomarkers, providing potential diagnostic or therapeutic targets. In summary, we provide a roadmap of endothelial cell identity in normal and malignant brain tissue, using a bioinformatics-based method developed to resolve bulk RNAseq datasets into constituent cell type-enriched profiles.

## INTRODUCTION

Comprehensive characterisation of human organs, and their constitutive cell types, is required to fully understand biological processes and disease development; concepts underlying large-scale tissue and cell profiling projects, such as the Human Protein Atlas (Uhlen et al., 2015) and Human Cell Atlas (Regev et al., 2017). RNAseq data from unfractionated human normal and diseased tissue is freely accessible through online portals, e.g. Genotype-tissue expression (GTEx) project (https://gtexportal.org) (Consortium, 2015) and the Cancer Genome Atlas (https://cancergenome.nih.gov), but deciphering cell type-specific profiles in such mixed-cell ‘bulk’ data is difficult. Recent advances have facilitated sequencing of discrete cell populations or individual cells, but practical and technical challenges, such as sourcing of suitable material, occurrence of artefacts due to processing, compromised read depth and financial constraints, limit the accessibility of such methods (Beliakova-Bethell et al., 2014; Rizzetto et al., 2017; Saliba et al., 2014; Ziegenhain et al., 2017). At the individual cell-type level, one of the most well-studied organs is the brain; a complex structure composed primarily of neurons, together with astrocytes, oligodendrocytes, microglia and an extensive vascular network of endothelial cells (EC) and associated mural cells (Azevedo et al., 2009; von Bartheld et al., 2016; Zhao et al., 2015). Previous studies have isolated and analysed the transcriptomes of these brain cell types, primarily from mouse (La Manno et al., 2018; Pandey et al., 2018; Saunders et al., 2018; Vanlandewijck et al., 2018; Zeisel et al., 2015), but also human tissue (Darmanis et al., 2015; Reddy et al., 2017; Zhang et al., 2016). The most common primary brain malignancy is glioma, of which 60-70% of cases are the most advanced form, glioblastoma multiforme (GBM), an incurable disease with a median survival of 15 months (Lim et al., 2018; Osuka and Van Meir, 2017). Cross-talk between EC and tumour cells is important for GBM progression (Yan et al., 2017) and studies indicate that GBM EC can be derived from the tumour cells themselves (Zhao et al., 2018). Bulk sequencing approaches have identified GBM molecular signatures (Jovcevska, 2018), but do not resolve disease-associated cell-type specific changes. Few studies have analysed GBM on a cell-by-cell basis, and it remains challenging to identify cell-type specific modifications, particularly of those cells present in relatively low abundance, such as EC, due to the small number of tumours analysed (Darmanis et al., 2017; Patel et al., 2014; Yuan et al., 2018).

Previously, we identified the core *body-wide* EC-enriched transcriptome from unfractionated RNAseq from 32 different organs (Butler et al., 2016), using a bioinformatics-analysis based on the relative proportion of EC across samples. Here, we analysed unfractionated RNAseq from samples of a *single* organ type, the human cortex, to identify endothelial, astrocyte, oligodendrocyte and neuron cell type-enriched genes. We used RNAseq from 516 lower grade gliomas (LGG) and 401 GBM to decipher global EC-compartment modifications, identifying tumour specific EC biomarkers. We provide a web-based tool to explore cell-type expression profiles in normal and malignant brain (https://cellcorrelation.shinyapps.io/Brain/ [*working version URL*]). Our approach has comparable reliability with isolated cell RNAseq for the identification of highly enriched cell type genes and does not require high-level bioinformatics expertise. It can be used to analyse and compare existing RNAseq datasets, regardless of processing or analysis platforms, to extract new biological insights into health and disease.

## RESULTS

### Cell type reference transcripts correlate across unfractionated cortex RNAseq data

RNAseq data from unfractionated normal brain samples (cortex region) was sourced from (i) GTEx Analysis V7 (n=158 samples) (https://gtexportal.org) (Consortium, 2015) and (ii) AMP-AD knowledge portal (MAYO RNAseq study, controls [n=80]) (https://www.synapse.org) (Allen et al., 2016). We selected ‘reference’ gene transcripts, which encode for established cell-type specific markers for: (**A**) Endothelial cells (EC): [*CD34, CLEC14A, VWF*] (**B**) Astrocytes (AC): [*BMPR1B, AQP4, SOX9*] (**C**) Oligodendrocytes (OC): [*MOG, CNP, MAG*] and (**D**) Neurons (NC): [*TMEM130, STMN2, THY1*] (Butler et al., 2016; Cahoy et al., 2008; Darmanis et al., 2015; Pfeiffer et al., 1993; Sun et al., 2017). We analysed GTEx cortex RNAseq data to calculate correlation coefficient values (corr.) between these cell-type reference transcripts across samples. Consistent with co-expression, reference transcripts within each cell-type group correlated with each other: EC [*CD34, CLEC14A, VWF*] mean corr. 0.66 p>0.0001, AC [*BMPR1B, AQP4, SOX9*] mean corr. 0.81 p>0.0001, OC [*MOG, CNP, MAG*] mean corr. 0.92 p>0.0001, NC [*TMEM130, STMN2, THY1*] corr. 0.89 p>0.0001, whilst reference transcripts between cell-type groups did not (Figure S1).

### Reference transcript analysis can resolve cell type genes from cortex RNAseq data

We performed a full analysis of the GTEx cortex RNAseq data to produce correlation values between each reference transcript and the other >20,000 mapped protein coding genes. A high mean correlation coefficient with the cell-type reference transcripts should indicate enrichment of the gene(s) in question in that cell type. To test sensitivity and specificity of the method, we compared correlation coefficients between the reference transcripts and 6 ‘*test*-panels’ - genes categorised in the literature as enriched in: (1) brain EC [*test*-EC] (2) AC [*test*-AC] (3) OC [*test*-OC] (4) NC [*test*-NC] (5) microglia [*test*-MG] and (6) smooth muscle [*test*-SMC] (Cahoy et al., 2008; Chu and Peters, 2008; Conley, 2001; Darmanis et al., 2015; Dreiza et al., 2010; He et al., 2016; Long et al., 2009; Miwa et al., 1991; Rensen et al., 2007; Wang et al., 2003; Yamawaki et al., 2001; Zhang et al., 2014) (Supplemental Table 1, Tabs 1-4: Column A). Each set of reference transcripts correlated most highly with genes in the corresponding *test*-panel (Figure 1A), with no overlap with genes from any other *test*-panel (Supplemental Table 1, Tabs 1-4: Column A). See Supplemental Materials page 14 for further details. We performed an equivalent *test*-panel analysis using RNAseq data from the AMP-AD knowledge portal (MAYO RNAseq study, control cortex samples, n=80). Correlation coefficients between the *test*-panels and the EC, AC, OC and NC reference transcripts in the MAYO (Figure 1B i-iv, respectively), were comparable to the GTEx (Table S1, Tab 1-4). For *test*-EC, AC, OC and NC genes, correlation values vs. the corresponding reference transcripts were high in both GTEx and MAYO data; the resultant cluster lying in the upper quadrant of the comparative plot (Figure 1B); the shaded box and dashed lines indicate the selected threshold requirement for classification as cell-type enriched (Figure S2). *Test*-MG and *test*-SMC genes did not highly correlate with the EC, AC, OC or NC reference transcripts (Figure 1 Ai-iv). However, to further verify that MG and SMC-enriched transcripts would not be incorrectly classified as EC-, AC-, OC-or NC-enriched, we selected 3 known MG [*C1QA, AIF1, LAPTM5*] (Fonseca et al., 2017; Ito et al., 1998; Zhang et al., 2014) and SMC [*FHL5, ACTA2, ACTG2*] (Halim et al., 2017; Vanlandewijck et al., 2018) transcripts and calculated their correlation with the *test*-panels (GTEx) (Table S1, Tab 5). All *test*-transcripts correlated more highly with the corresponding reference transcripts, than with MG or SMC reference transcripts (Figure S3 Ai-iv). Higher correlation values of the MG-or SMC-reference transcripts with the *test*-EC panel (vs. *test*-AC, OC or NC), indicated that MG or SMC genes were most likely to be incorrectly classified as EC-enriched (Figure S3 Ai), rather than AC-, OC-or NC-enriched. Transcripts identified as EC-, AC-, OC-or NC-enriched were excluded if the mean correlation with the reference transcripts < mean corr. vs. MG or SMC transcripts (GTEx vs. GTEx data). As predicted, most exclusions were from the EC-enriched list (Figure S2).

**TABLE 1.**
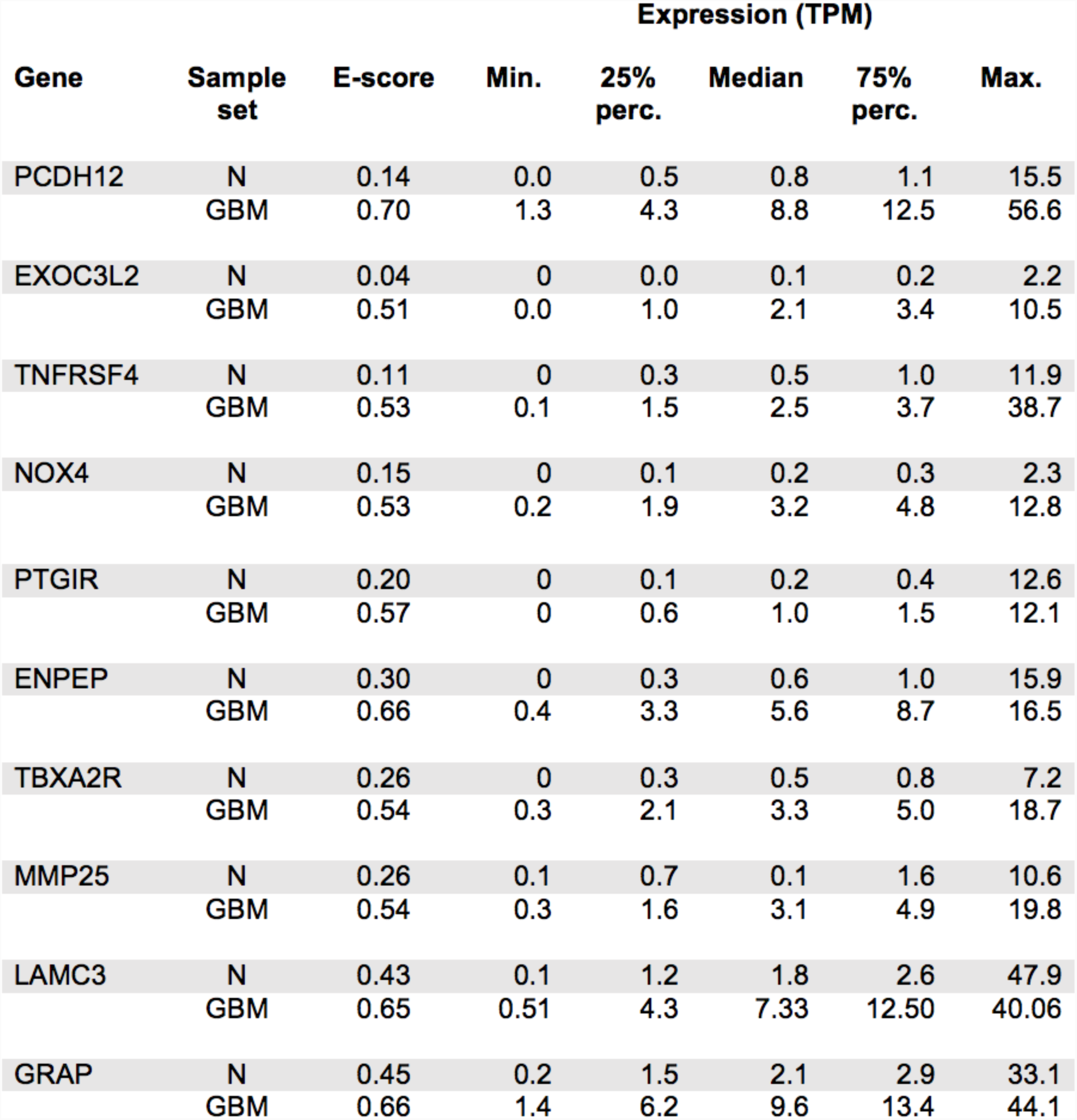
Glioblastoma multiforme (GBM) specific EC-enriched genes. Identified GBM EC markers (’Gene’) and corresponding comparative expression data downloaded from the OASIS portal (Fernandez-Banet et al., 2016). Sample set: **N** - normal brain tissue (n=424), **GBM** - glioblastoma multiforme (n=169), **E-score** ‘enrichment score’ - mean correlation coefficient with EC reference transcripts [*CD34, CLEC14A, VWF*].

| Gene | Sample set | E-score | Expression (TPM) |  |  |  |  |
| --- | --- | --- | --- | --- | --- | --- | --- |
|  |  |  | Min. | 25% perc. | Median | 75% perc. | Max. |
| PCDH12 | N | 0.14 | 0.0 | 0.5 | 0.8 | 1.1 | 15.5 |
|  | GBM | 0.70 | 1.3 | 4.3 | 8.8 | 12.5 | 56.6 |
| EXOC3L2 | N | 0.04 | 0 | 0.0 | 0.1 | 0.2 | 2.2 |
|  | GBM | 0.51 | 0.0 | 1.0 | 2.1 | 3.4 | 10.5 |
| TNFRSF4 | N | 0.11 | 0 | 0.3 | 0.5 | 1.0 | 11.9 |
|  | GBM | 0.53 | 0.1 | 1.5 | 2.5 | 3.7 | 38.7 |
| NOX4 | N | 0.15 | 0 | 0.1 | 0.2 | 0.3 | 2.3 |
|  | GBM | 0.53 | 0.2 | 1.9 | 3.2 | 4.8 | 12.8 |
| PTGIR | N | 0.20 | 0 | 0.1 | 0.2 | 0.4 | 12.6 |
|  | GBM | 0.57 | 0 | 0.6 | 1.0 | 1.5 | 12.1 |
| ENPEP | N | 0.30 | 0 | 0.3 | 0.6 | 1.0 | 15.9 |
|  | GBM | 0.66 | 0.4 | 3.3 | 5.6 | 8.7 | 16.5 |
| TBXA2R | N | 0.26 | 0 | 0.3 | 0.5 | 0.8 | 7.2 |
|  | GBM | 0.54 | 0.3 | 2.1 | 3.3 | 5.0 | 18.7 |
| MMP25 | N | 0.26 | 0.1 | 0.7 | 0.1 | 1.6 | 10.6 |
|  | GBM | 0.54 | 0.3 | 1.6 | 3.1 | 4.9 | 19.8 |
| LAMC3 | N | 0.43 | 0.1 | 1.2 | 1.8 | 2.6 | 47.9 |
|  | GBM | 0.65 | 0.51 | 4.3 | 7.33 | 12.50 | 40.06 |
| GRAP | N | 0.45 | 0.2 | 1.5 | 2.1 | 2.9 | 33.1 |
|  | GBM | 0.66 | 1.4 | 6.2 | 9.6 | 13.4 | 44.1 |

**FIGURE 1.**
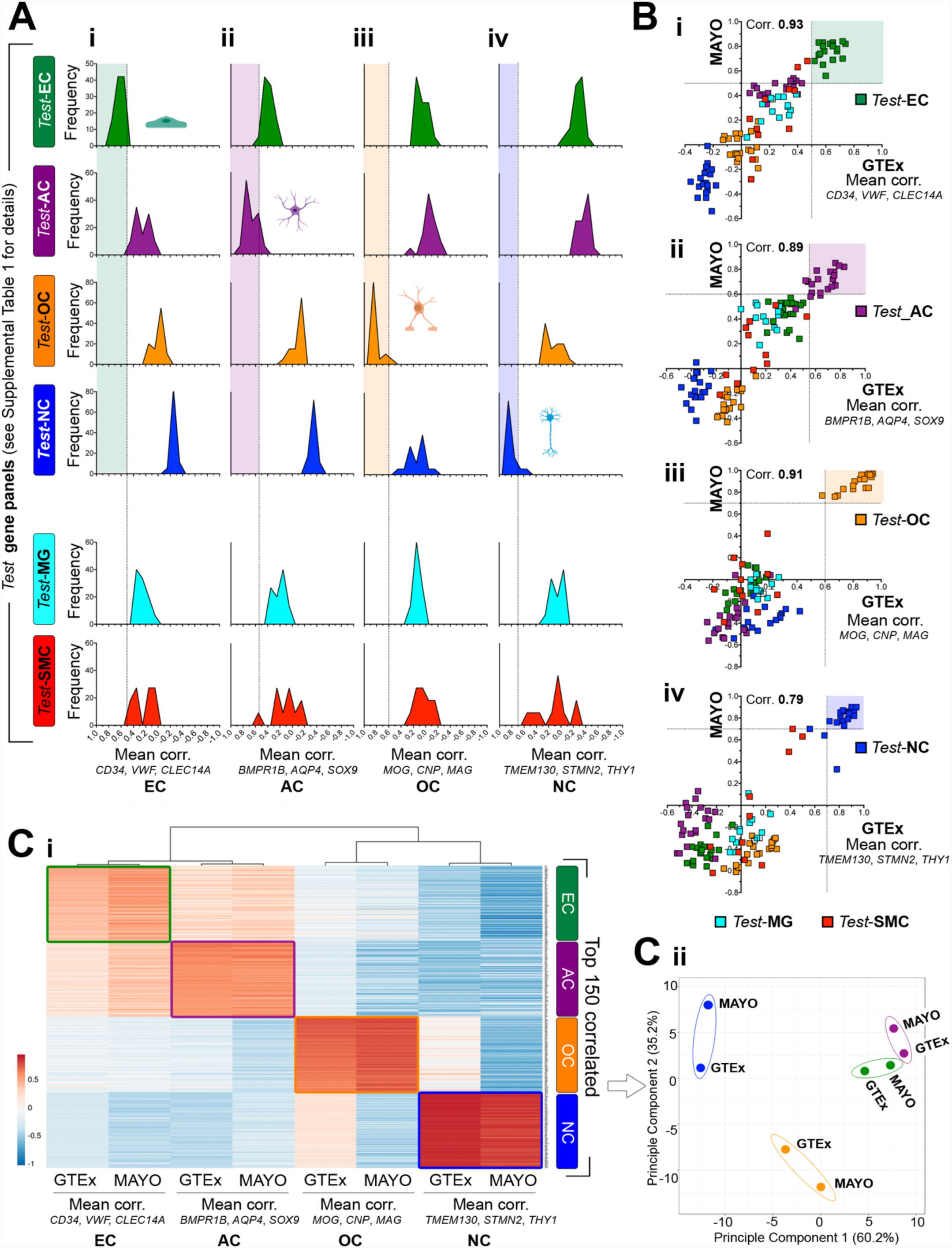
Reference transcript based analysis identifies cell type-enriched genes from unfractionated human cortex data. RNAseq data from unfractionated normal human cortex from the Genotype-Tissue Expression [GTEx] portal (n=158) was used to calculate Spearman’s correlation coefficients between reference transcripts selected for: (**A**) (i) endothelial cells (EC) [*CD34, CLEC14A, VWF*], (ii) astrocytes (AC) [*BMPR1B, AQP4, SOX9*], (iii) oligodendrocytes (OC) [*MOG, CNP, MAG*], (iv) neurons (NC) [*TMEM130, STMN2, THY1*] and ‘*test*-panels’ - genes previously categorised in the literature as enriched in: brain EC [*test*-EC], AC [*test*-AC], OC [*test*-OC], NC [*test*-NC], microglia [*test*-MG] or smooth muscle cells [*test*-SMC]. Frequency distribution of coefficient vales are plotted and shaded areas indicate thresholds selected. (**B**) Equivalent *test*-panel analysis was performed on RNAseq data from unfractionated normal human cortex samples from the AMP-AD knowledge portal MAYO RNAseq study (controls, n=80) and means compared to the GTEX results for (i) EC, (ii) AC, (iii) OC and (iv) NC reference transcripts. Dotted lines indicate the dual-thresholding criteria applied for classification as cell-type enriched. (**C**) Heat map plot of correlation values of the top 150 genes identified as EC-, AC-, OC-or NC-enriched vs. all the reference transcript sets, and (**D**) corresponding principle component analysis. Adapted from images generated by ClustVis (<u>/clustvis/</u>). See also Table S1-5.

### EC and AC have a panel of dual-enriched transcripts

We compared the relationships between transcripts in the cell-type enriched lists. To compare groups, e.g. EC-vs. AC-enriched, the following values were calculated for each transcript featuring in either list: (1) the difference between the mean correlation coefficients for both sets of corresponding reference transcripts i.e. EC: [*CD34, CLEC14A, VWF*] and AC: [*BMPR1B, AQP4, SOX9*] (the ‘differential correlation score’) and (2) the ‘enrichment ranking’, based on the correlation value with each set of reference transcripts (highest correlation: rank=1) (Figure S3B). Threshold lines indicate the rank number below which transcripts were classified as cell-type enriched. EC and AC-enriched transcripts ‘crossed’ on the plot (Figure S3 Bi, yellow points), whilst EC transcripts well separated from the enriched gene profiles of the other cell types (Figure S3 Bii and iii). OC-and NC-enriched transcripts were also well separated (Figure S3 Biv). 31 transcripts fulfilled the necessary criteria to be classified both as EC *and* AC enriched (Figure 2A, yellow points [plot as for Figure S3B, but transcripts ranked below enriched classification threshold excluded]) (Table S3, Tab 5). However, the majority of transcripts in the EC-and AC-enriched lists were well separated by differential correlation. We sourced data from a previously published study where human brain tissue was sorted into cell populations prior to RNAseq (Zhang et al., 2016); for transcripts that we identified as (i) AC-enriched *only*, (ii) AC *and* EC-enriched or (iii) EC-enriched *only (*Figure 2A: purple, yellow and green, respectively) expression profiles were consistent with our classifications (Figure 2B). Based on these analyses, transcripts classified as EC-or AC-enriched were excluded from the final list(s) if they fulfilled enrichment criteria for both, or if they had higher correlation values with the other (non-corresponding) reference transcripts.

**FIGURE 2.**
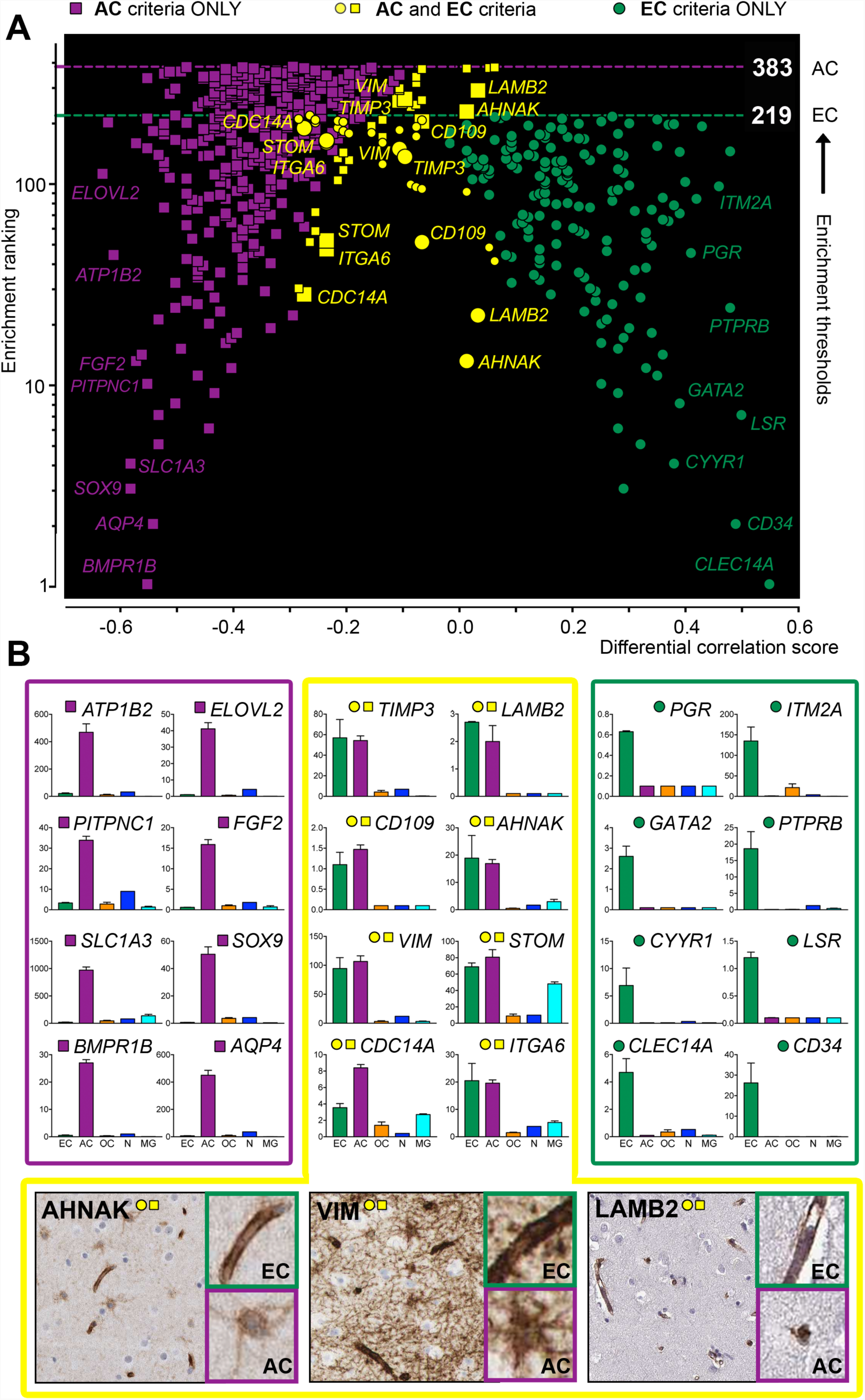
Reference transcript based analysis identifies dual cell type-enriched genes from unfractionated human cortex data. RNAseq data from unfractionated normal human cortex from the Genotype-Tissue Expression [GTEx] portal (n=158) was used to calculate Spearman’s correlation coefficients between the reference transcripts for endothelial cells (EC) [*CD34, CLEC14A, VWF*] or astrocytes (AC) [*BMPR1B, AQP4, SOX9*]. (**A)** For transcripts fulfilling criteria for classification as EC-(circles) or AC-enriched (squares), the ‘*differential correlation score’ (*difference between mean corr. with EC and AC reference transcripts) was plotted vs. ‘enrichment ranking’ (position in each respective enriched list, highest correlation = ranking 1). Threshold lines denote ranking below which transcripts were classified as EC-or AC-enriched. Yellow symbols represent transcripts classified as *both* EC-and AC-enriched (circular *and* square symbol, on the same X-axis dimension). (**B)** mRNA expression data from 5 brain cell types: EC (green), AC (purple), oligodendrocyte, OC (orange), neurons, NC (blue) and microglia, MG (cyan), isolated from human brain prior to RNAseq (Zhang et al., 2016), for genes we classified as AC-enriched (purple box), dual-enriched in AC and EC (yellow box) and EC-enriched (green). Protein profiling of human cortex tissue for expression of *AHNAK, VIM* and *LAMB2* in both AC and EC. See also Table S3, Tab 5.

### Identification of endothelial, astrocyte, oligodendrocyte and neuron-enriched genes

Following application of selection criteria (reference transcript correlation in GTEx, replication in MAYO, FDR threshold, SMC/MG and dual enriched transcript exclusion) (Figure S2), 166 genes were classified as EC-enriched, 351 were AC-enriched, 380 were OC-enriched and 2015 were NC-enriched (Table S2-5 [enriched: Tab 1, all values: Tab 2]). Expression of selected transcripts was confirmed in the expected cell type by immunohistochemistry (Figure S4). A heat map plot of the correlation values of the top 150 genes identified as EC-, AC-, OC-or NC-enriched vs. the EC, AC, OC and NC reference genes, respectively, revealed the resolution of cell-type expression profiles in the GTEx and MAYO datasets (Figure 1Ci). Principle component analysis revealed the MAYO and GTEx data for each cell category clustered together, and EC-and AC-enriched genes clustered more closely than any of the other cell types (Figure 1Cii). The top 15 most highly enriched genes for each category contained known cell-enriched transcripts (Figure 3, marked in bold). Gene Ontology (GO) and Reactome (‘pathway’) analysis (Ashburner et al., 2000) was performed on the final list of EC-, AC-, OC-and NC-enriched transcripts. The most significant biological process GO groups in the EC-enriched list were related to EC function, including ‘*vasculature development*’ and ‘*angiogenesis*’ (p-values <3.8×10^17) and reactome pathways included ‘*haemostasis*’ and ‘*NOSTRIN mediated eNOS trafficking*’ [p-values <3.9×10^5] (Table S2, Tab 4). The most significant biological process GO groups in the AC-enriched list included ‘*regulation of signalling*’ and ‘*small molecule catabolic process*’ (p-values <1.4×10^10) and reactome pathways included ‘*metabolism*’ and ‘*transport of small molecules*’ (p-values <1.3×10^5) (Table S3, Tab 4). The most significant biological process GO group in the OC-enriched list was ‘*myelination*’ (p-value <1.6×10^17) and the single reactome pathway identified was ‘*transport of small molecules*’ (p-value <7.8×10^7) (Table S4, Tab 4). The most significant biological process GO groups in the NC-enriched list included ‘*nervous system development*’ and ‘*trans-synaptic signalling*’ (p-values <2.2×10^32) and reactome pathways included ‘*neuronal system*’ and ‘*neurotransmitter receptors and postsynaptic signal transmission*’ (p-values <9.5×10^17) (Table S3, Tab 4). Summary plots were generated using REViGO (Supek et al., 2011) (Figure 3).

**FIGURE 3.**
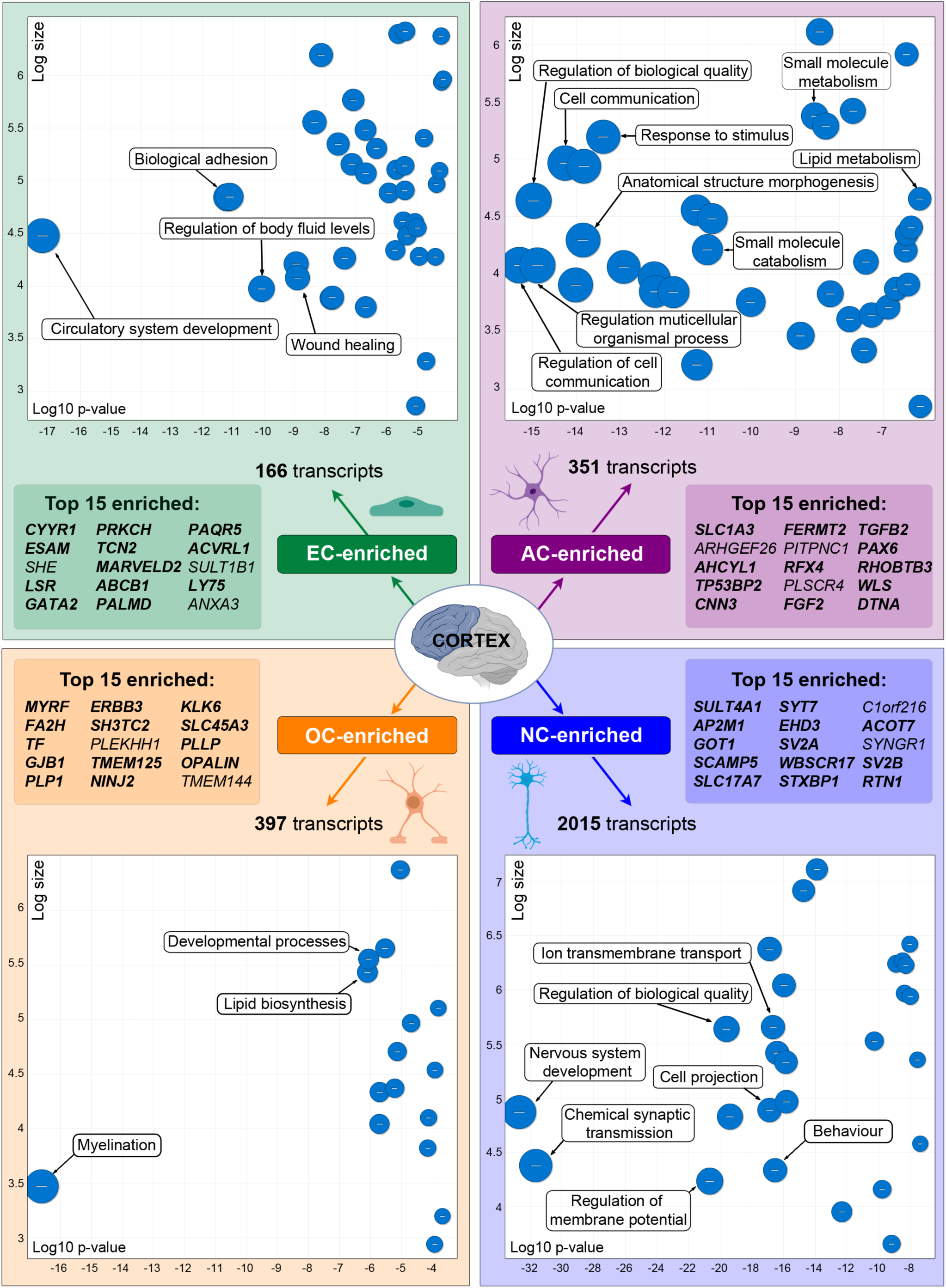
Transcripts classified as endothelial (EC)-, astrocyte (AC)-, oligodendrocyte (OC)-or neuron (NC)-enriched in human brain cortex. Human cortex RNAseq from the Genotype-Tissue Expression [GTEx] portal (n=158), and the AMP-AD knowledge portal MAYO RNAseq study (n=80) was used to identify EC-, AC-, OC-and NC-enriched transcripts. Total enriched transcript number, top 15 most enriched (bold denotes previously described) and a summary of gene ontology (GO) enrichment for all enriched genes in each cell type (generated using REViGO) are presented. See also Table S2-S5 and Figure S2.

### Comparison of results with single-cell or fractionated brain cell type RNAseq data

We compared the top 100 most enriched EC, AC, OC and NC transcripts from our unfractionated (UF) RNAseq analysis, to existing single-cell sequencing (scRNAseq) of human brain (H1) (Darmanis et al., 2015) and RNAseq of fractionated FACS sorted or immuno-panned isolated cell populations from both human (H2) (Zhang et al., 2016) and mouse (M1) brain (Zhang et al., 2014). The top 100 most enriched genes we identified in the EC, AC, OC and NC categories were significantly enriched in the corresponding human single cell type sequencing analysis (H1): total with ROTS score (see methods for ROTS definition) >2 vs. other cell types EC 76/100, AC 94/100, OC 91/100, NC 98/100 (Figure 4Ai and ii) and the human fractionated cell type analysis (H2): fold expression >2 vs. other cell types EC 84/100, AC 90/100, OC 95/100, NC 85/100 (all p>0.001 vs. other cell types) (Figure 4Bi and ii). Agreement was less consistent in the mouse fractionated cell type analysis (M1): fold expression >2 vs. other cell types EC 74/100, AC 56/100, OC 57/100, NC 65/100 (p>0.0001 vs. other cell types) (Figure 4Ci and ii), consistent with differences in gene expression and cell type specificity between human and murine brain. Overall, ~50% of the top 100 EC-, AC-, OC-and NC-enriched genes were categorised as enriched in the same cell type in *all* 3 (H1, H2 and M1) datasets (Figure 4Di-iv), and ≥98% were enriched in *at least* one other dataset. Thus, our results were consistent with fractionated tissue or scRNAseq based cell-type transcript classification.

**FIGURE 4.**
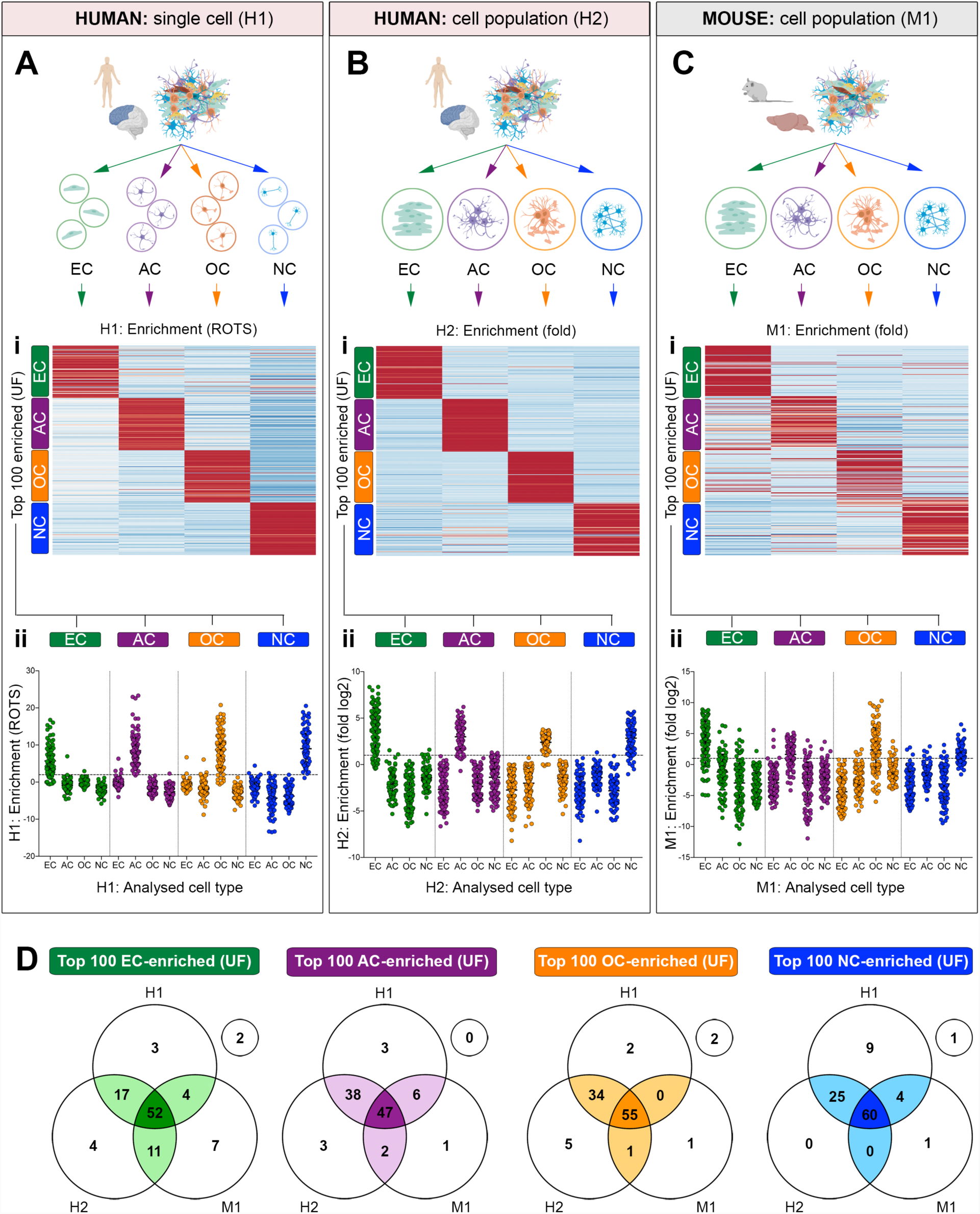
Comparison of transcripts classified as endothelial (EC)-, astrocyte (AC)-, oligodendrocyte (OD)-or neuron (NC)-enriched with existing single-cell or fractionated brain cell type RNAseq data. Datasets from (**A)** single-cell sequencing of human brain (scRNAseq) (Darmanis et al., 2015) (H1) or RNAseq of isolated cell populations from (**B)** human (H2) (Zhang et al., 2016) or (**C)** mouse (M1) brain (Zhang et al., 2014) were downloaded from the respective publications. Transcript ‘enrichment scores’ in EC, AC, OC and NC populations in H1 (‘ROTS score’), H2 and M2 (fold enrichment vs. other cell types) were compiled for the top 100 transcripts identified as EC-, AC-, OC-or NC-enriched from our unfractionated (UF) RNAseq analysis and presented as: (**i**) heat maps and (**ii**) grouped plots. Horizontal line = threshold for ‘enrichment’. (**D)** Venn diagrams showing number of top 100 transcripts we identified as EC-, AC-, OC-or NC-enriched that were also identified as enriched in the corresponding cell type in H1, H2, M1 datasets (free circles – UF only).

### Comparison of reference transcript-based and weighted correlation network analysis

A disadvantage of our analysis is its reliance on a pre-selected panel of only 3 cell type-enriched ‘reference’ transcripts, and thus, is subject to a possible input bias. To determine the extent of this limitation, we analysed the same datasets using an alternative ‘unbiased’ approach - weighted correlation network analysis (‘WGCNA’) (Langfelder and Horvath, 2008); where correlations were generated between each transcript and all others, and transcripts with similar profiles were clustered together. Analysis of the GTEx and MAYO datasets produced a total of 37 and 50 independent clusters, respectively (Table S6, Tab 1, annotated with arbitrary numbers). *Test*-OC and *test*-EC panels were used to identify clusters representing these cell types (Table S6, Tab 2) (Figure S5 A and B, green/orange boxes) and GO enrichment analysis of these clusters were consistent with OC or EC identify, respectively (Figure S5C). The majority of genes classified as OC-or EC-enriched by the reference transcript analysis were also classified as such by WGCNA (Figure S5 Ei and Fii) (337/397 [85%] and 106/166 [64%], respectively). See Supplemental Materials page 15-17 for details. We compared EC and OC-enriched lists, generated using each method, to brain cortex single cell RNAseq (Darmanis et al., 2015) [H1] and isolated brain cell type RNAseq (Zhang et al., 2016) [H2] (see Figure 4). The majority of the OC-enriched genes identified by the reference transcript or WGCNA, were correspondingly enriched in both the [H1] (ref. transcript 82% vs. WGCNA 76%) (Figure S5 Gi) and [H2] (ref. transcript 80% vs. WGCNA 78%) datasets (Figure S5 Gii). Similarly, the majority of EC-enriched genes identified by the reference transcript or WGCNA, were correspondingly enriched in both the [H1] dataset (ref. transcript 76% vs. WGCNA 72%) (Figure 5Hi) and the [H2] dataset (ref. transcript 83% vs. WGCNA 67%) datasets (Figure S5 Hii). Overall, fewer genes were classified as OC-or EC-enriched by the reference transcript analysis (OC: 379 vs. 446, and EC: 166 vs. 252), but results were better supported by data from [H1] and [H2], compared to the WGCNA.

**FIGURE 5.**
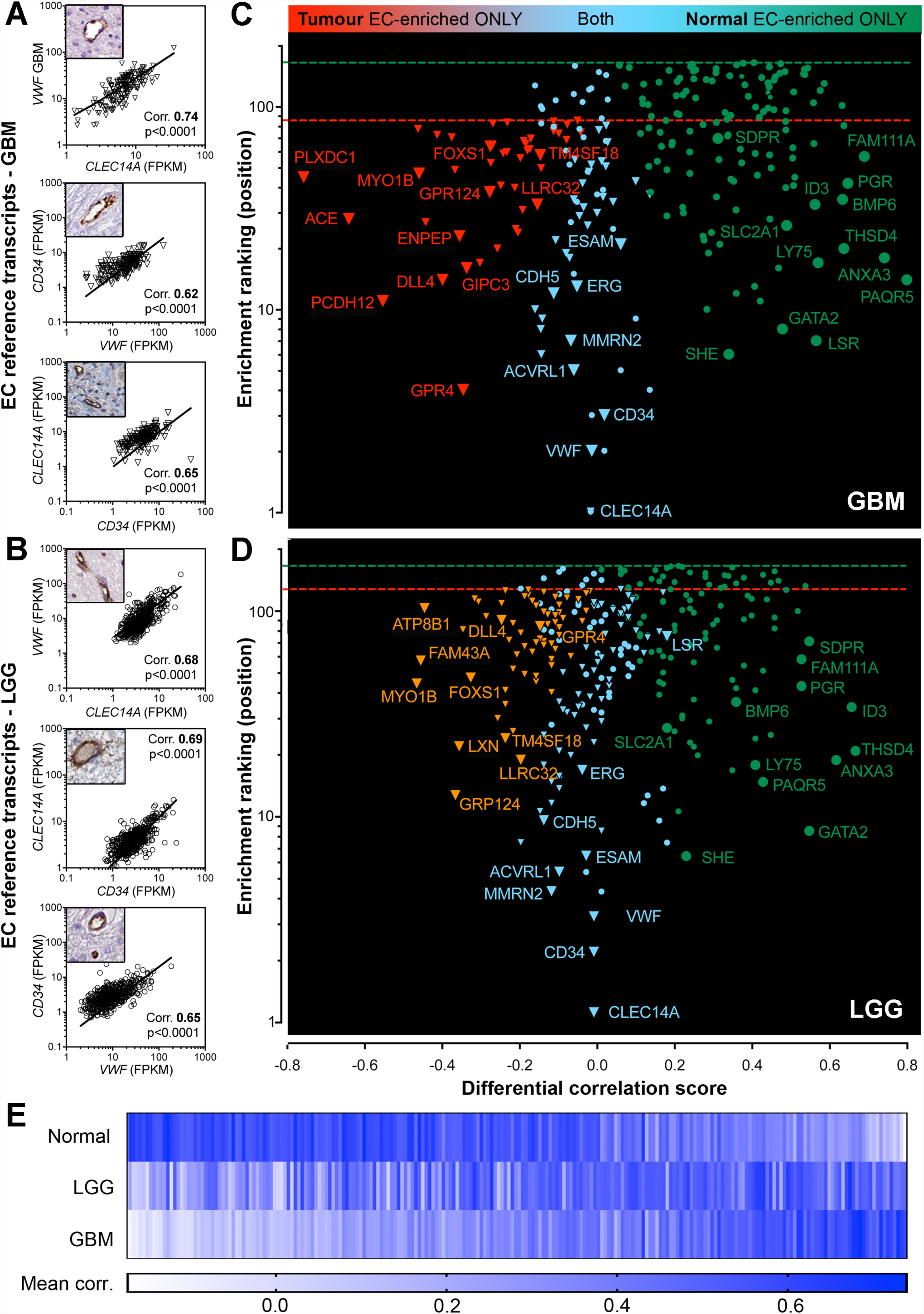
Reference transcript based analysis can resolve global modifications of EC-enriched transcriptome of glioblastoma multiforme and lower grade glioma. RNAseq data from unfractionated human glioblastoma multiforme (GBM) (n=401) and human lower grade glioma (LGG) (n=516) was downloaded from the Cancer Genome Atlas (TCGA) and Spearman’s correlation coefficients were calculated between the endothelial (EC) reference transcripts [*CD34, CLEC14A, VWF*] and all mapped protein-coding genes. Correlations between *CD34, CLEC14A, VWF* and IHC confirming EC restricted expression (label as for y-axis) in (**A)** GBM and (**B)** LGG. Plots showing *differential correlation score* and *enrichment rank* of transcripts categorised as EC-enriched in: (**C)** GBM and/or normal brain and (**D)** LGG and/or normal brain. (**E)** Heat map of correlation values between transcripts classified as EC-enriched in GBM and/or LGG and/or normal brain, and the EC reference transcripts. See also Table S7.

### Reference transcript analysis can be used to profile disease-associated modifications

Our data from the analysis of normal cortex was used as a starting point for the identification of cellular changes in malignant disease. We used the reference transcript analysis to study disease-associated changes in the vasculature of glioblastoma multiforme (GBM) and lower grade glioma (LGG), where cross-talk between EC and tumour cells promotes proliferation and subsequent disease progression (Yan et al., 2017). Expression tables for GBM (n=401) and LGG (n=516) samples from the Cancer Genome Atlas (TCGA) were sourced from the Genomic Data Commons Data Portal (https://portal.gdc.cancer.gov/). As in normal brain, EC genes [*CD34, CLEC14A, VWF*] correlated highly in GBM (Figure 5A) and LGG (Figure 5B) (corr. mean GBM 0.67, LGG 0.74), with EC restricted expression (Figure 5A and B, upper left images), indicating suitability as EC reference transcripts in these cohorts. GBM and LGG datasets were analysed as for normal brain (GTEx) (Table S7, Tab 1) to identify EC-enriched transcripts. WGCNA of GBM RNAseq produced 2 clusters which contained EC markers (n=104 and n=35) (Supplemental Figure 6A). These clusters contained 78/85 (92%) of genes classified as GBM EC-enriched by the reference transcript analysis (Supplemental Figure 6B) (Table S7, Tab 2). We compared GBM and normal brain, using the differential correlation score and enrichment rank of all transcripts categorised as EC-enriched in GBM *or* normal brain (Figure 5C). 33 transcripts were EC-enriched in normal brain *and* GBM (Figure 6C, blue data points) (Table S7, Tab 3). Of these, we previously identified 29/33 (88%) as body-wide core EC-enriched (Butler et al., 2016), including *ENG, TIE1, ESAM, CDH5* and *ERG (*which encodes for a key transcription factor in the maintenance of EC-identify (Shah et al., 2016)); thus, this group likely represents genes indispensable for vascular function. A panel of genes were EC-enriched in GBM, but not in normal brain, including *PLXDC1, ACE, PCDH12,* and vice versa, including *PAQR5, ANXA3, FAM111A (*Figure 6C, red and green data points, respectively) (Table S7, Tab 3). LGG had a higher proportion of EC-enriched genes in common with normal brain, compared to GBM (Figure 5D - more blue data points vs. Figure 5C) and correspondingly, fewer genes were modified in LGG vs. normal. Of those EC-enriched genes that were modified in LGG, parallels were observed with GBM e.g. *MYO1B, DLL4, FOXS1, GPR4, LLRC32, TM4SF18* ‘gained’ EC-enrichment and *SDPR, FAM111A, PGR, ANXA3, GATA2* ‘lost’ EC-enrichment, in both tumour grades (Figure 5C and D). A heat map plot of the raw correlation values, between the EC reference transcripts and those classified as EC-enriched in normal *or* LGG *or* GBM (Figure 5E), was consistent with the hypothesis that LGG represents an ‘intermediate state’ between normal and GBM.

**FIGURE 6.**
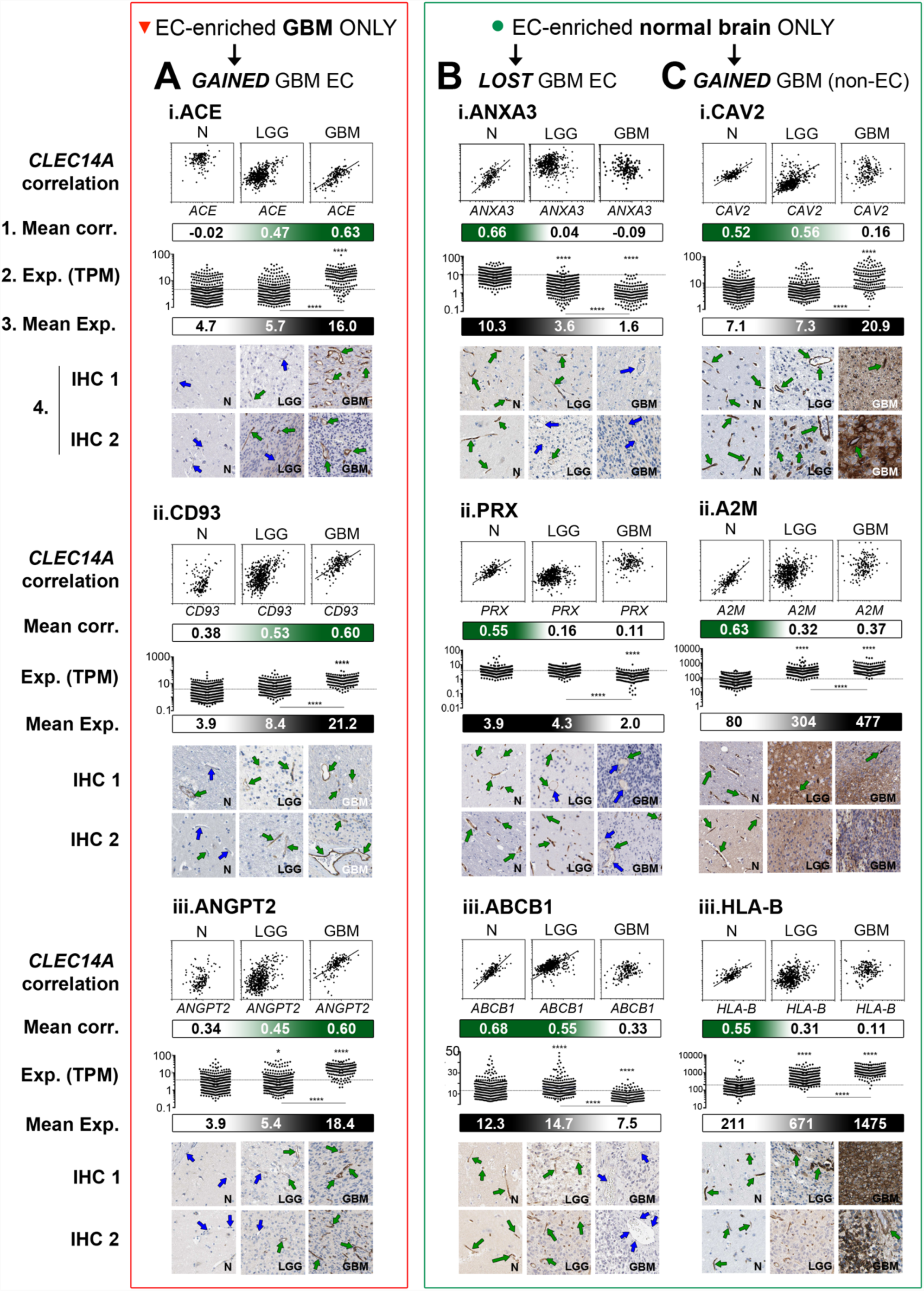
Using reference transcript based correlation analysis with mRNA expression and tissue protein profiling to decipher nature of EC-enriched modifications. RNAseq data from unfractionated normal human cortex (n=158) (N), human lower grade glioma (LGG) (n=516) and human glioblastoma multiforme (n=401) (GBM) was used to calculate: (**1**) mean Spearman’s correlation coefficient between endothelial (EC) reference transcripts [*CD34, CLEC14A, VWF*] and selected genes (upper plot shows *CLEC14A* vs. gene). mRNA expression vales (TPM) for selected genes were downloaded from the OASIS portal (Fernandez-Banet et al., 2016) for N, LGG and GBM and displayed as (**2**) individual samples and (**3**) as a mean expression in all. (**4**) Protein profiling was performed for selected genes on tissue sections of N, LGG and GBM. Candidate genes were classified as: (**A)** EC-enriched in glioblastoma multiforme (GBM), but not normal (N) tissue (red box) ([**i**] ACE, [**ii**] CD93 and [**iii**] ANGPT2], EC-enriched in normal (N) tissue, but not GBM (green box) either due to: (**B)** *loss* of expression from GBM EC ([i] *ANXA3*, [ii] *PRX* and [iii] *ABCB1*], or (**C)** gain of expression outside the vasculature in GBM ([i] CAV2, [ii] A2M and [iii] HLA-B] *p<0.05, ****p<0.0001. See also Table S7.

### Further resolution of GBM-associated EC-enriched transcriptome modifications using mRNA expression data in combination with protein profiling

We selected a panel of EC genes identified as modified in GBM vs. normal brain to further characterise, by combining expression data (normal, LGG and GBM) from the OASIS portal (Fernandez-Banet et al., 2016) with protein profiling, which we performed as part of the Human Protein Atlas (HPA) Pathology Atlas (Uhlen et al., 2017). ACE (angiotensin converting enzyme) was classified as EC-enriched in GBM, but not normal brain (mean corr. 0.63 GBM vs. −0.02 normal) correspondingly, *ACE* mRNA was higher in GBM vs. normal (mean TPM 16.0 GBM vs. 4.7 normal p<0.0001) (Figure 6Ai, exp.(TPM)). The ‘degree’ of *ACE* EC-enrichment and expression was intermediate in LGG, between normal and GBM (Figure 6Ai), indicating changes occurred in LGG, but they were not as pronounced (or consistent) as those in the higher grade GBM. ACE protein was not detected in normal tissue (Figure 6Ai, IHC 1 and 2) (blue arrows indicate negative vessels), was variable in LGG EC, and was highly abundant in GBM EC (green arrows indicate positive staining). Similar patterns were observed for CD93 (receptor for the C1q complement factor) (Figure 6Aii) and ANGPT2 (angiopoietin 2) (Figure 6Aiii).

When transcripts were classified as EC-enriched in normal brain, but not in GBM (Figure 5C, green data points), this could indicate that: (1) expression was *lost* from GBM EC, or (2) expression was *gained* outside the vasculature i.e. in tumour tissue (illustrated in Figure 7A and B). *ANXA3 (*Annexin 3A) was EC-enriched in normal brain, but not in GBM (mean corr. 0.66 normal vs. −0.09 GBM) and *ANXA3* mRNA was decreased in GBM vs. normal (mean TPM 1.6 GBM vs. 10.3 normal p<0.0001) (Figure 6Bi). IHC showed ANXA3 was expressed in normal EC, was variable in LGG EC, and was absent from GBM EC. Similar patterns were observed for *PRX (*Periaxin) (Figure 6Bii) and *ABCB1 (*ATP-binding cassette sub-family B member 1) (Figure 6Biii). *CAV2 (*Caveolin 2) was also EC-enriched in normal brain, but not GBM (mean corr. 0.52 normal vs. 0.16 GBM) (Figure 6Ci) however, *CAV2* mRNA was *increased* in GBM vs. normal (mean TPM 20.9 GBM vs. 7.1 normal p<0.0001), and CAV2 was expressed outside the EC compartment (Figure 6Ci). Similar was observed for *A2M (*Figure 6Cii) and *HLA-B (*Figure 6Ciii). Generally, LGG represented an ‘intermediate state’ between normal brain and GBM, the degree of similarity to either condition depending on the specific transcript.

**FIGURE 7.**
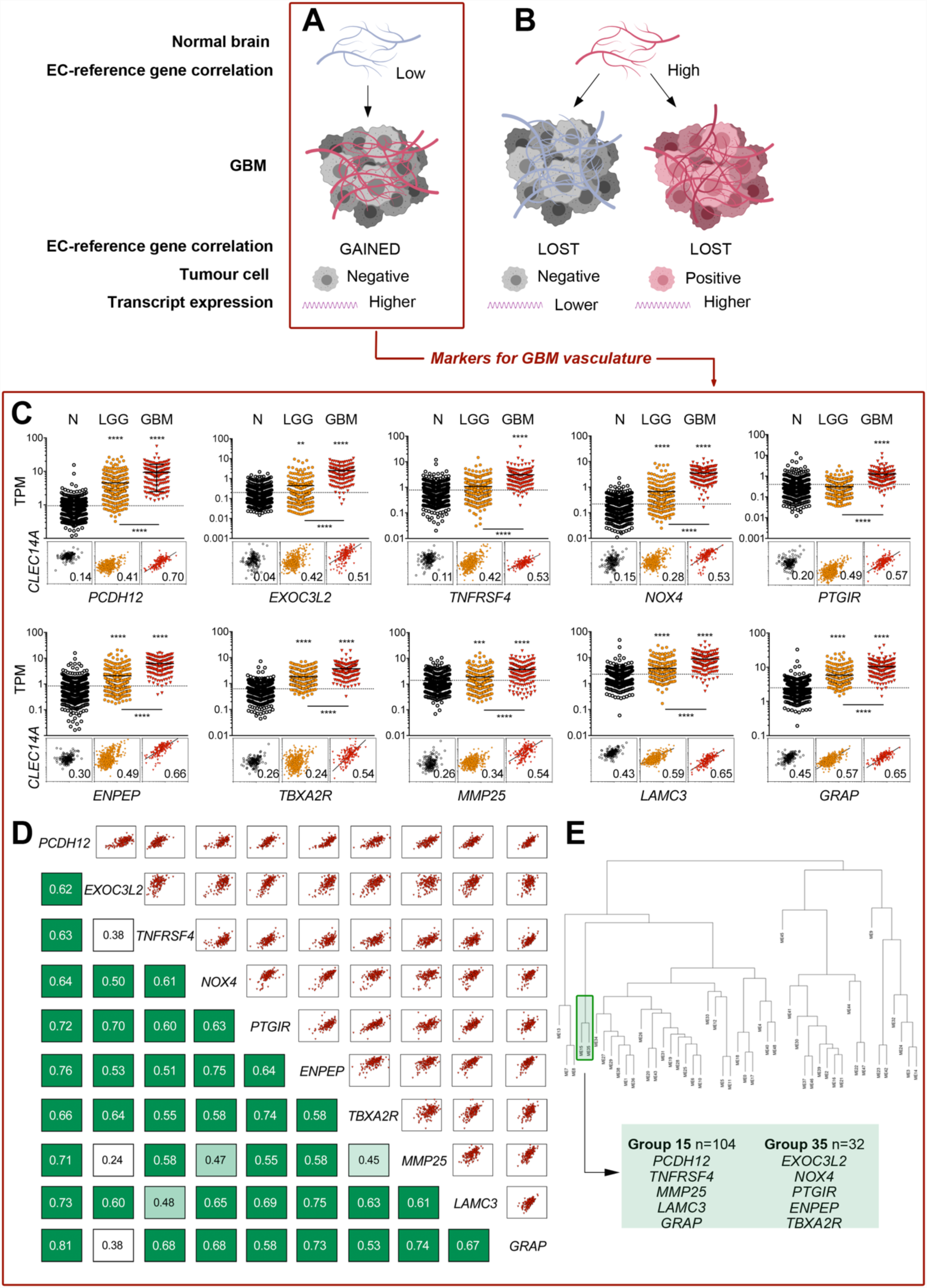
Glioblastoma multiforme (GBM) specific EC-enriched genes. Summary of normal brain EC-enriched transcriptome modifications associated with GBM: (**A)** EC-transcript expression specific to GBM only (*increased* EC-enrichment score), (**B)** EC-transcript expression absent in GBM EC *or* gained in non-EC compartment (both *decreased* EC-enrichment score). (**C)** mRNA expression data (TPM) for GBM EC marker candidates in normal brain (N), lower grade glioma (LGG) and GBM and illustrative correlation plots vs. EC reference transcript *CLEC14A,* with mean values vs. all EC-reference transcripts [*CD34, CLEC14A, VWF*] in bottom right of plot. (D) Correlation matrix of GBM marker candidates generated from GBM RNAseq samples (E) Weighted correlation network analysis (WGCNA) dendrogram of unfractionated GBM: EC groups identified by green boxes ** p<0.01, ***p<0.001 ****p<0.0001. See also Table S7.

We identified 10 GBM EC markers (Figure 7C), based on the following criteria: [1] they were *not* expressed in EC from multiple other vascular beds (Butler et al., 2016) [2] they had *low* expression levels in the normal brain (mean TPM<5) [3] they had a high differential EC-enriched correlation between GBM and normal (Table 1). The 10 GBM-EC markers correlated with each other in the GBM RNAseq (Figure 7D), but not the normal RNASeq (mean corr. GBM 0.61 ± 0.06 vs. normal 0.18 ± 0.05). Consistent with previous observations, LGG represented an intermediate between GBM and normal (mean corr. LGG 0.46 ± 0.04) (correlation matrices, Table S7, Tab 4). All 10 GBM-EC markers were also in the EC-annotated groups in the WGCNA of the GBM RNAseq (Figure 7E, Figure S6), consistent with co-expression in GBM EC. Thus, our method can be applied to perform a systems-based identification of highly GBM-specific EC-markers, that could have clinical applications for tumour grading, prognosis prediction and therapeutic targetting.

## DISCUSSION

We have demonstrated that publically available expression data can be used, in combination with our method, to provide a road map of cell type-enriched transcripts in the human brain, and to provide an unprecedented resolution of the changes in the EC-enriched transcriptome associated with glioblastoma multiforme (GBM) and lower grade glioma (LGG). We provide searchable tables (Table S2-7) and a web interface (https://cellcorrelation.shinyapps.io/Brain/ [*working version URL*]) for users to determine the extent of endothelial, astrocyte-, oligodendrocyte-or neuron-specificity of any mapped protein coding gene in the normal cortex and EC enrichment score in GBM or LGG.

Our approach has some advantages over other types of transcriptomic profiling, such as single cell sequencing (scRNAseq). scRNAseq typically involves the extraction and sequencing of thousands of individual cells (Darmanis et al. 2015; Villani et al. 2017; Pandey et al. 2018; Papalexi and Satija 2018) to detect cell type specific genes and identify to rare cell subtypes in heterogeneous tissue. In contrast, our analysis does not require removal of cells from their resident niche, circumventing modifications due to loss of environmental cues, or processing-associated activation (Chow and Gu, 2015; Urich et al., 2012; Wilhelm et al., 2016) whilst avoiding the introduction of technical variability (Beliakova-Bethell et al., 2014; Saliba et al., 2014). As sample collection and processing needs to be tightly controlled, many brain scRNAseq studies use non-human tissue, e.g. (Pandey et al., 2018; Rosenberg et al., 2018; Vanlandewijck et al., 2018). In contrast, our method can be used to analyse hundreds of human biological replicates concurrently, extracting information from bulk RNAseq data, from normal or diseased tissue. Results can be directly compared across datasets, regardless of processing or analysis platforms, at no experimental expense. However, our analysis does have limitations; we applied strict thresholds to maximise confidence in our classifications and to only identify the most highly specific genes for each cell type. It was not feasible to identify an extensive panel of transcripts expressed at moderately higher levels in one cell type vs. others, which is reflected by the comparatively low number of genes we annotated as cell-enriched, compared to isolated brain cell type or scRNAseq studies (Darmanis et al., 2015; Vanlandewijck et al., 2018; Zhang et al., 2014). A further limitation is the possible incorrect classification of transcripts, particularly between closely associated cell types, which could have comparable ratios across samples, such as EC with pericytes or smooth muscle cells (SMC). We used test-panels to exclude SMC as a source of false positive EC-enriched genes, but equivalent analysis for pericytes was more challenging, due to a lack of specific markers (Armulik et al., 2011). However, genes previously described as pericyte-enriched, e.g. PDGFRB, ACTA2, DES, MCAM (Smyth et al., 2018), VTN and IFITM1 (He et al., 2016), were not classified as EC-enriched. ANPEP (CD13), a marker often used to identify pericytes in mouse (Crouch and Doetsch, 2018), was classified as brain EC-enriched in our study, however, it was also classified as such by scRNAseq (Darmanis et al., 2015) and is highly expressed in EC lines (Thul et al., 2017). Thus, whilst our data indicates that pericyte genes are not generally incorrectly classified as EC-enriched, we cannot definitively exclude the possibility that the EC-enriched list contains dual pericyte/EC-enriched transcripts.

We performed a global analysis to identify changes in the brain EC-enriched transcriptome in GBM. Bulk RNAseq has been used to define GBM molecular signatures (Jovcevska, 2018), but not to resolve cell-type specific changes. A small number of studies have dissociated GBM tissue for the analysis of gene expression on a cell-by-cell basis (Darmanis et al., 2017; Dieterich et al., 2012; Patel et al., 2014; Yuan et al., 2018), but it remains challenging to define GBM-specific cell type signatures, particularly of minority cells, such as EC, due to the small number of tumours analysed. Brain EC are highly specialised with extensive adherens and tight junctions (Tietz and Engelhardt, 2015) and reduced pinocytosis, transcytosis, non-selective fenestration and leukocyte adhesion molecule expression (Zhao et al., 2015). We identified transcripts enriched in normal brain EC, but not in GBM EC, including those encoding for specialised brain EC proteins, such as the drug efflux pump TP-binding cassette sub-family B member 1 (ABCB1) (Borst and Schinkel, 2013), and the immuno-and permeability regulatory protein Periaxin (PRX) (Wang et al., 2018). As specialised brain EC features are induced and regulated by the local microenvironment (Chow and Gu, 2015) and rapidly lost ex vivo (Urich et al., 2012; Wilhelm et al., 2011) one could speculate that GBM EC fail to express such genes due to the loss of the normal microenvironment. The majority of transcripts we identified as having EC-specific expression in both normal and GBM were ‘core’ EC genes, found throughout human tissue beds (Butler et al., 2016); suggesting GBM EC maintain critical genes for basic identify and function.

We identified a panel of GBM EC markers, some of which have been previously reported, including CD93 (Dieterich et al., 2012; Langenkamp et al., 2015) and ANGPT2 (Dieterich et al., 2012; Scholz et al., 2016; Stratmann et al., 1998). The majority, to our knowledge, were previously unknown, e.g. exocyst complex component 3-like 2 (EXOC3L2), TNF receptor superfamily member 4 (TNFRSF4), Thromboxane A2 Receptor (TBXA2R) and Prostaglandin I2 Receptor (PTGIR). EXOC3L2, a component of the exocyst involved in vesicle fusion with the plasma membrane, is poorly characterised; the only published study on the protein describes its upregulation in mouse embryonic sprouting blood vessels and induction in primary human cells in response to VEGF (Barkefors et al., 2011). This is consistent with its expression in GBM EC; GBM tumour cells produce VEGF, which induces angiogenesis (Das and Marsden, 2013), however, the role of EXOC3L2, or potential as angiogenic target in GBM remains unknown. TNFRSF4 (Ox40) has been a target of experimental tumour immunotherapy, including in glioma, due to its expression on tumour infiltrating T-cells (Buchan et al., 2018). In a mouse sarcoma model, TNFRSF4 was also expressed on EC which, following TNFRSF4-agonist treatment, responded with (T-cell independent) VCAM-1 expression (Pardee et al., 2010). Thus, one could speculate that TNFRSF4 expression on GBM EC could indicate a role for a direct EC-interaction in the antitumor effect of TNFRSF4-targeted immunotherapy (Curry et al., 2016). TBXA2R and PTGIR encode for prostaglandin receptors, which bind thromboxane A2 and prostacyclin, respectively, products of the short lived intermediate molecule prostaglandin H_2_, generated from arachidonic acid, via cyclooxygenase (COX)-1 or −2. Several studies have demonstrated that COX-2 is elevated in GBM, and levels positively correlated with tumour grade, reoccurrence and shorter survival (Qiu et al., 2017), however, the expression of prostaglandin receptors on GBM EC, and the functional consequences, have not been explored. A panel of GBM EC-enriched genes we identified are linked through the renin-angiotensin system (RAS): angiotensin I converting enzyme (ACE), glutamyl aminopeptidase (ENPEP) and NADPH oxidase 4 (NOX4). RAS inhibitors suppress progression and lengthen survival period in several cancer types (Ishikane and Takahashi-Yanaga, 2018; Pinter and Jain, 2017), including glioma (Levin et al., 2017), where they have a broad spectrum effectiveness, and are currently being considered for inclusion in treatment protocols (Perdomo-Pantoja et al., 2018). ACE and ENPEP are central enzymes in the RAS, catalysing the conversion of angiotensin I to angiotensin II, and angiotensin II to angiotensin III, respectively. Angiotensin II and III have various effects, mediated primarily through angiotensin receptor I (ATR1) (Jackson et al., 2018), such as the induction of pro-survival signalling, via NF-KB mediated anti-apoptotic molecules, or through PI3K-Akt mediated suppression of caspases (George et al., 2010), the induction of cellular proliferation (Clark et al., 2011; Perdomo-Pantoja et al., 2018) and loss of blood brain EC barrier properties (Biancardi and Stern, 2016). More high grade astrocytoma tumour cells express ATR1, compared to lower grade, and expression is associated with cell proliferation, vascularisation and shorter survival (Arrieta et al., 2008). Angiotensin II induces NOX4, which regulates the production of reactive oxygen species (ROS) (Nguyen Dinh Cat et al., 2013). ROS play an important role in signal transduction, cell differentiation, tumour cell proliferation, apoptosis and angiogenesis (Guo and Chen, 2015). NOX4 is more highly expressed in GBM than lower grade tumours, with a role in tumour proliferation and resistance to apoptosis induced by chemotherapeutic agents (Shono et al., 2008). Despite the acknowledged importance of RAS in cancer, the specific cell types underlying the increased signalling are not well described. Based on our data, we speculate that GBM EC drive increased local levels of angiotensin II and II, which promote cancer progression and maintenance.

In summary, we identify cell-enriched genes from unfractionated brain RNAseq, without the need for high-level bioinformatics expertise or complex modelling. We model system-level changes in the EC-compartment associated with brain malignancy of increasing severity, providing biological insight and identifying potential targets for therapy.

## METHODS AND RESOURCES

### CONTACT FOR REAGENT AND RESOURCE SHARING

Dr. Lynn Marie Butler

Clinical Chemistry and Blood Coagulation, Department of Molecular Medicine and Surgery, Karolinska Institute, SE-171 76 Stockholm, Sweden

### EXPERIMENTAL MODEL AND SUBJECT DETAILS

Human tissue protein profiling was performed in house as part of the Human Protein Atlas (HPA) project (Ponten et al., 2008; Uhlen et al., 2015; Uhlen et al., 2017) (www.proteinatlas.org). Normal brain (cortex), lower grade glioma (LGG) and glioblastoma multiforme (GBM) samples were obtained from the Department of Pathology, Uppsala University Hospital, Uppsala, Sweden; as part of the Uppsala Biobank. Samples were handled in accordance with Swedish laws and regulations, with approval from the Uppsala Ethical Review Board (Uhlen et al., 2015).

Bulk RNAseq data analysed in this study was part of the Genotype-Tissue Expression (GTEx) Project (gtexportal.org) (Consortium, 2015) (dbGaP Accession phs000424.v7.p2), the TCGA Research Network (cancergenome.nih.gov) and AMP-AD Knowledge Portal (www.synapse.org). Datasets from normal brain scRNAseq (Darmanis et al., 2015) and RNAseq of isolated human and mouse brain cell types (Zhang et al., 2014; Zhang et al., 2016) were downloaded from the Gene Expression Omnibus (GEO) database (Accession IDs: GSE67835, GSE73721 and GSE52564, respectively). Comparative transcript expression between normal, LGG and GBM samples were downloaded from the OASIS portal (oasis-genomics.org) (Fernandez-Banet et al., 2016).

## EXPERIMENTAL METHODS

### Tissue Profiling: Human tissue sections

Tissue sections from human cerebral cortex were generated and stained, as previously described (Ponten et al., 2008; Uhlen et al., 2015). Briefly, formalin fixed and paraffin embedded tissue samples were sectioned, de-paraffinised in xylene, hydrated in graded alcohols and blocked for endogenous peroxidase in 0.3% hydrogen peroxide diluted in 95% ethanol. For antigen retrieval, a Decloaking chamber® (Biocare Medical, CA) was used. Slides were boiled in Citrate buffer®, pH6 (Lab Vision, CA). Primary antibodies and a dextran polymer visualization system (UltraVision LP HRP polymer®, Lab Vision) were incubated for 30 min each at room temperature and slides were developed for 10 minutes using Diaminobenzidine (Lab Vision) as the chromogen. Slides were counterstained in Mayers hematoxylin (Histolab) and scanned using Scanscope XT (Aperio). Primary antibodies, source, target and identifier: Atlas Antibodies: AHNAK (HPA026643), A2M (HPA002265), ALDH1L1 (HPA036900), ANPEP (HPA004625), ANXA3 (HPA013398), CALD1 (HPA008066), CARNS1 (HPA038569), CD34 (HPA036722), CD93 (HPA012368), CLDN11 (HPA013166), CLEC14A (HPA039468), DLG3 (HPA001733), GGT5 (HPA008121), GJA1 (HPA035097), HSDL2 (HPA050453), ITGB1 (HPA059297), KANK3 (HPA051153), KLC1 (HPA052450), MAP2 (HPA008273), MCC (HPA037391), MOAP1 (HPA000939), NRGN (HPA038171), PAK1 (HPA003565), PHLDB1 (HPA037959), PLP1 (HPA004128), PROCR (HPA039461), PRSS23 (HPA030591), PRX (HPA001868), PSMB9 (HPA053280), TAGLN2 (HPA001925), TINAGL1 (HPA048695), TLN1 (HPA004748), VWF (HPA001815). Abnova: DCN (H00001634-M01). Agilent: VIM (M7020). Leica Biosystems: MBP (NCL-MBP), ACE (NCL-CD143-510). Merck: (ABCB1 (MAB4120), FGF2 (AB1458), SLC2A1 (AB1341). R&D Systems: PON2 (AF4344). Santa Cruz Biotechnology: ANGPT2 (sc-74403), HLA-B (sc-55582), LIPE (sc-74489), PGM5 (sc-73613). Sigma-Aldrich: LAMB2 (AMAb91097). Thermo Fisher Scientific: CAV2 (41-0700), CLDN5 (RB-9243).

## QUANTIFICATION AND STATISTICAL ANALYSIS

### Reference transcript based correlation analysis

This method was adapted from that previously developed to determine the cross-tissue pan-EC-enriched transcriptome (Butler et al., 2016). As different cell types are present in different proportions across individual samples, we used a correlation analysis to identify cell-type specific transcripts. We calculated the pairwise Spearman correlation coefficients between reference transcripts selected as proxy markers for: endothelial cells (EC): [*CD34, CLEC14A, VWF*] (in GTEx, TCGA [LGG and GBM] and MAYO datasets), astrocytes (AC): [*BMPR1B, AQP4, SOX9*] (in GTEx and MAYO datasets), oligodendrocytes (OC) [*MOG, CNP, MAG*] (GTEx and MAYO datasets), neurons (NC) the genes [*TMEM130, STMN2, THY1*] (GTEx and MAYO datasets) and all mapped protein coding genes. To exclude false positives, we also calculated correlations between proxy markers for microglial cells (MG) [*C1QA, AIF1, LAPTM5*] and SMC [*FHL5, ACTA2, ACTG2*] and selected transcripts identified as cell-type enriched. Reference transcripts used in analysis of **GTEx**: EC, AC, OC, NC, MG, SMC; **LGG**: EC; **GBM**: EC; **MAYO**: EC, AC, OC, NC. Non-coding transcripts and transcripts with expression TPM <0.1 were excluded from final data tables. See results section for full analysis and exclusion criteria required for transcript classification as cell-type enriched. Correlation coefficients were calculated in R (v 3.4.3) using the *corr.test* function from the *psych* package (v 1.8.4). In addition to correlation coefficients False Discovery Rate (FDR) adjusted p-values (using Bonferroni correction) and raw p-values were calculated.

### Weighted correlation network (WGCNA) analysis

The R package WGCNA was used to perform co-expression network analysis for gene clustering, on log2 expression values. The analysis was performed according to recommendations in the WGCNA manual. Genes with too many missing values were excluded using the goodSamplesGenes() function. The remaining genes were used to cluster the samples, and obvious outlier samples were excluded. Using these genes and samples a soft-thresholding power was selected and the networks were constructed using a minimum module size of 15 and merging threshold of 0.05. Eigengenes were calculates from the resulting clusters and eigengene dendrograms were constructed using the plotEigengeneNetworks() function.

### Brain single-cell and isolated cell fraction RNAseq datasets

Brain cell type expression datasets from scRNAseq (Darmanis et al., 2015) (GSE67835) and isolated cell type RNAseq (Zhang et al., 2014; Zhang et al., 2016) (GSE73721 and GSE52564) studies were used to calculate cell type specific enrichment values for each mapped transcript. For scRNAseq cell-type enrichment was calculated using the ROTS analysis method (Suomi et al., 2017) and enrichment was defined as ROTS score >2. For the isolated cell type RNAseq fold-enrichment values in each cell type: EC, AC, OC (*myelinating oligodendrocytes* and *newly formed oligodendrocytes* were combined), NC and MG were calculated as described in the original studies; ‘*FPKM expression in one cell type divided by the average expression level in all other cell types*’. Cell enrichment was defined as a fold-enrichment of >2.

### Gene Ontology (GO) Enrichment analysis

The Gene Ontology Consortium (Ashburner et al., 2000) and PANTHER classification resource (Mi et al., 2013; Mi et al., 2016) were used to identify over represented terms (biological processes) in the panel of identified cell-type-enriched transcripts from the GO ontology database (release date March 2016).

## ADDITIONAL RESOURCES

The Human Protein Atlas (HPA) website contains details of all antibody-based protein profiling used in this study: www.proteinatlas.org. A searchable web-based interface can be used to explore the datasets presented here, on a transcript-by-transcript basis (https://cellcorrelation.shinyapps.io/Brain/ [working version URL])).

## AUTHOR CONTRIBUTIONS

Conceptualisation L.M.B.; Methodology P.D, B.H, L.M.B.; Formal analysis P.D, B.H, L.M.B.; Investigation P.D, L.M.B.; Resources T.R, M.U.; Writing – Original Draft P.D, L.M.B.; Writing – Review & Editing, All; Visualisation P.D, L.M.B.; Funding Acquisition J.O, L.M.B.

## ACKNOWLEDGEMENTS

Funding was granted to L.M.B. from Hjärt Lungfonden (20150623 and 20170759) and Vetenskapsrådet (2013-42608-102305-28). PD was supported by a postdoc grant from Strategic Research Areas (SFO), KTH to JO and LB. We also acknowledge funding from Stockholm County Council (LS 1302-0311) to JO. We acknowledge the staff of the Human Protein Atlas (HPA) program, the Science for Life Laboratory and the pathology team in Mumbai, India. We thank the Department of Pathology at the Uppsala Akademiska hospital, Uppsala, Sweden and Uppsala Biobank for kindly providing specimens used in this study. The HPA was funded by Knut & Alice Wallenberg Foundation.

## Data usage acknowledgements

The results shown here are based upon analysis of data generated as part of the Genotype-Tissue Expression (GTEx) Project (gtexportal.org) (Consortium, 2015), the TCGA Research Network (cancergenome.nih.gov) and AMP-AD Knowledge Portal (www.synapse.org). The GTEx project was supported by the Office of the Director of the National Institutes of Health, and by NCI, NHGRI, NHLBI, NIDA, NIMH, and NINDS. The Mayo RNAseq study data were provided by the following sources: The Mayo Clinic Alzheimers Disease Genetic Studies, led by Dr. Nilufer Taner and Dr. Steven G. Younkin, Mayo Clinic, Jacksonville, FL using samples from the Mayo Clinic Study of Aging, the Mayo Clinic Alzheimers Disease Research Center, and the Mayo Clinic Brain Bank. Data collection was supported through funding by NIA grants P50 AG016574, R01 AG032990, U01 AG046139, R01 AG018023, U01 AG006576, U01 AG006786, R01 AG025711, R01 AG017216, R01 AG003949, NINDS grant R01 NS080820, CurePSP Foundation, and support from Mayo Foundation. Study data includes samples collected through the Sun Health Research Institute Brain and Body Donation Program of Sun City, Arizona. The Brain and Body Donation Program is supported by the National Institute of Neurological Disorders and Stroke (U24 NS072026 National Brain and Tissue Resource for Parkinsons Disease and Related Disorders), the National Institute on Aging (P30 AG19610 Arizona Alzheimers Disease Core Center), the Arizona Department of Health Services (contract 211002, Arizona Alzheimers Research Center), the Arizona Biomedical Research Commission (contracts 4001, 0011, 05-901 and 1001 to the Arizona Parkinson’s Disease Consortium) and the Michael J. Fox Foundation for Parkinson’s Research.

