## Supplemental Data 1 for "A systems-based map of human brain cell-type enriched genes and malignancy-associated endothelial changes"

SUPPLEMENTAL FIGURE 1: **Reference transcript correlation matrix**

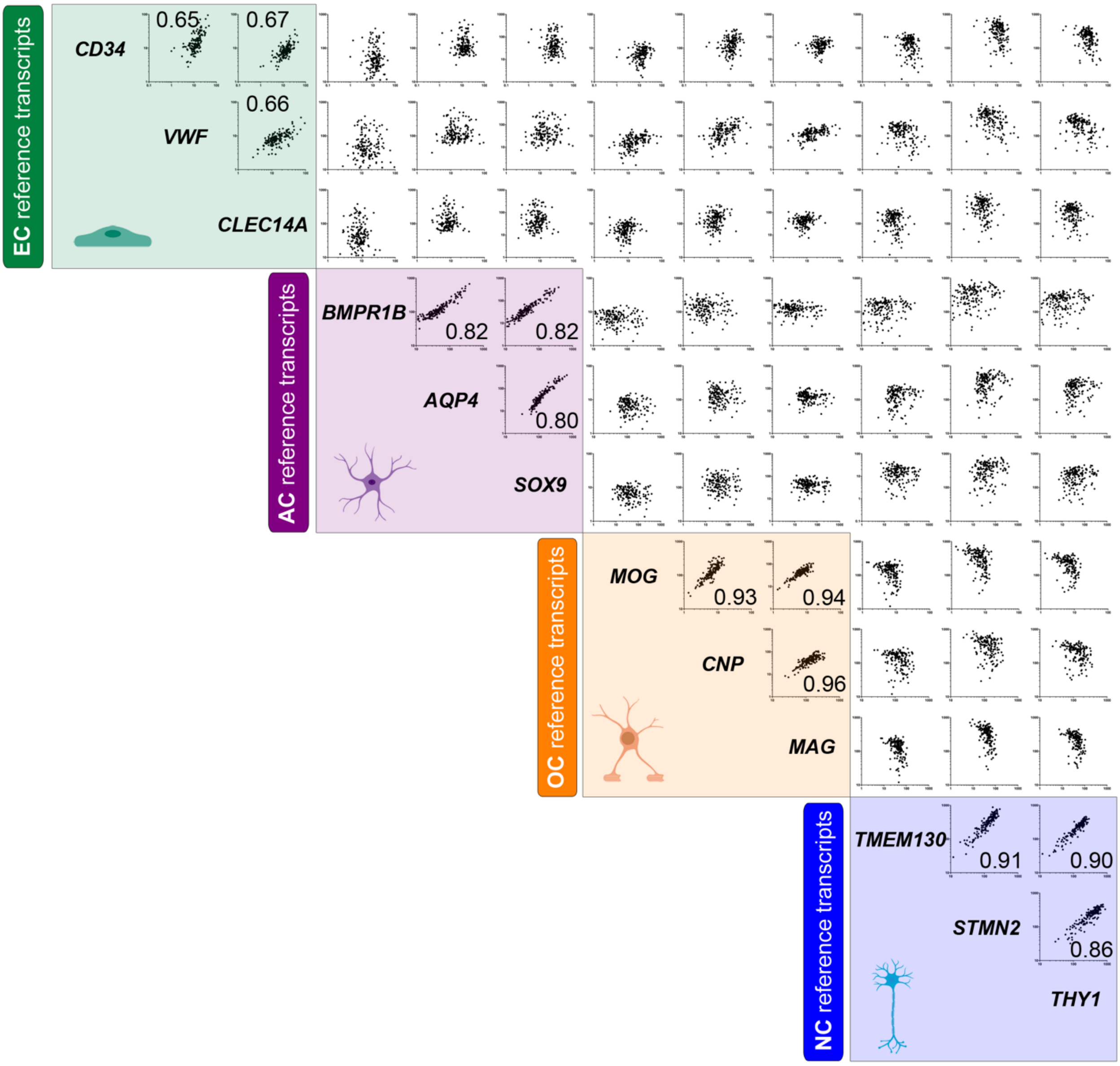


**FIGURE S1. Reference transcript correlation matrix: Related to Figure 1.** RNAseq data from unfractionated normal human cortex samples from the Genotype-Tissue Expression [GTEx] portal (n=158) was analysed to determine Spearman's correlation coefficients between 'reference' transcripts selected for endothelial cells (**EC**) [*CLEC14A, VWF*, *CD34*], astrocytes (**AC**) [*BMPR1B*, *AQP4*, *SOX9*], oligodendrocytes (**OC**) [*MOG*, *CNP*, *MAG*] and neurons (**NC**) [*TMEM130*, *STMN2*, *THY1*]. Plots show correlations between mRNA expression levels and numbers denote Spearman's correlation coefficients (all $p<0.0001$).

SUPPLEMENTAL FIGURE 2: **Cell type enrichment classification criteria**

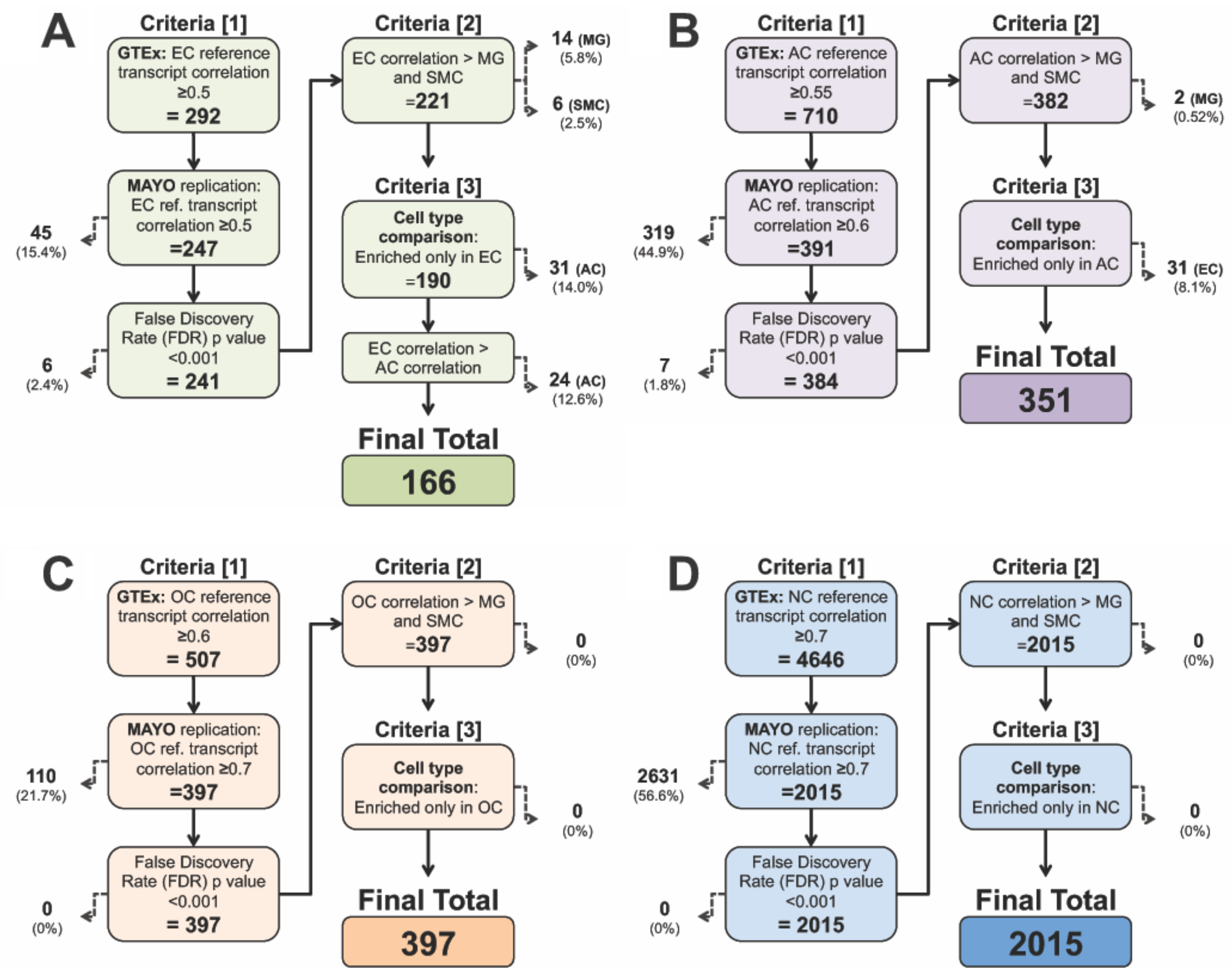


**FIGURE S2. Cell type enrichment classification criteria: Related to Figure 1-3.** Summary of analysis to identify (**A**) endothelial (EC)-, (**B**) astrocyte, (AC)- (**C**) oligodendrocyte (OC)- and (**D**) neuron (NC)-enriched transcripts using unfractionated brain cortex RNAseq data.

SUPPLEMENTAL FIGURE 3: **MG and SMC reference transcript correlations with *test*-panels and global relationships between cell-type enriched transcripts**

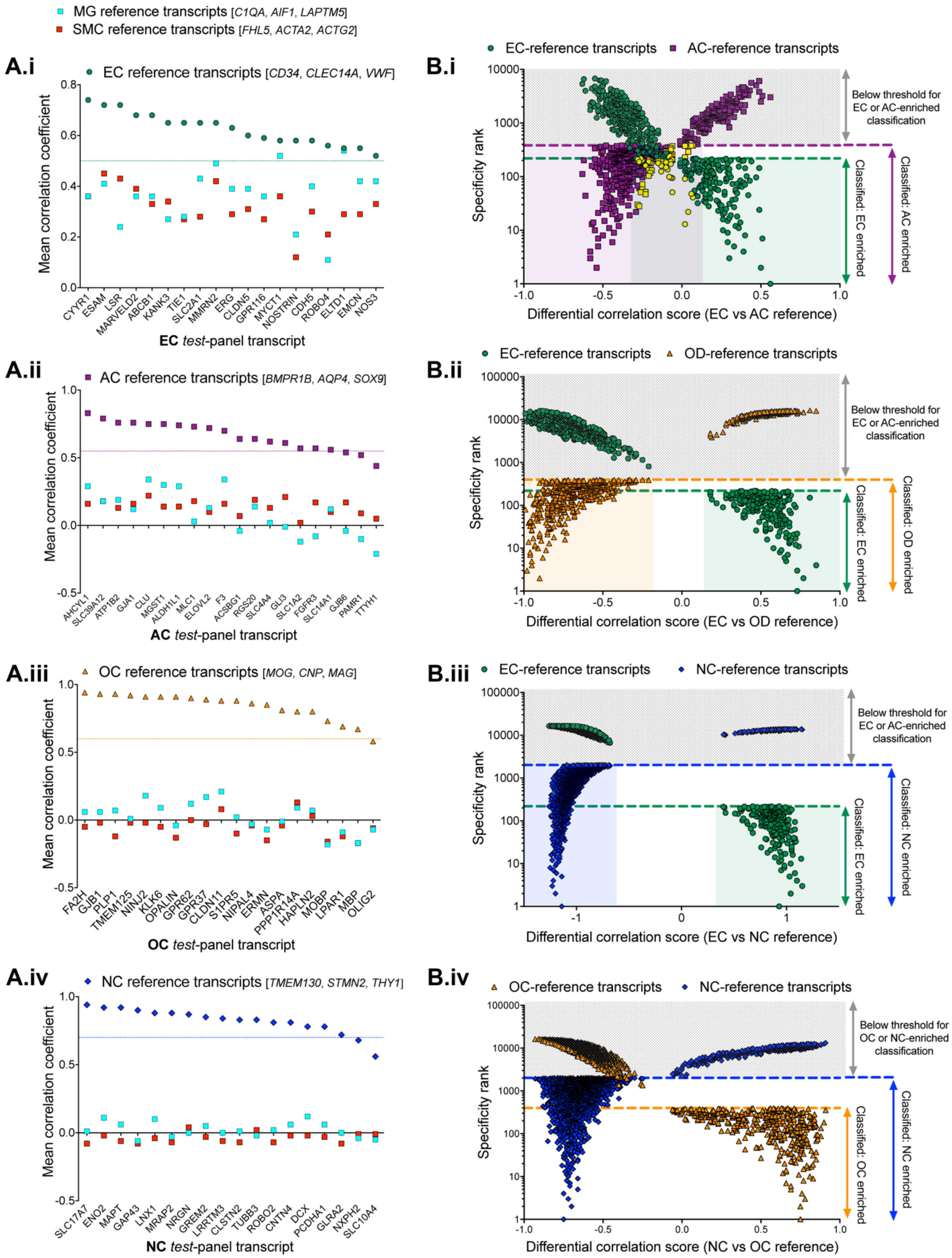

**FIGURE S3. Microglial and smooth muscle reference transcript correlations with *test*-panels and global relationships between cell-type enriched transcripts: Related to Figure 2 and 3. See also Table S1, Tab 5**. RNAseq data from unfractionated normal human cortex samples from the Genotype-Tissue Expression [GTEx] portal (n=158) was analysed to determine mean Spearman's correlation coefficients between microglial (MG) [*C1QA*, *AIF1*, *LAPTM5*] and smooth muscle cell (SMC) [*FHL5*, *ACTA2*, *ACTG2*] reference transcripts (mean corr. within each set: 0.94 and 0.72, respectively, both p<0.001) and genes in (**A**) (i) *test*-endothelial (EC), (**ii**) test-astrocyte (AC), (iii) *test*-oligodendrocyte (OC) and (iv) *test*-neuron NC panels. Correlation values for the corresponding reference transcripts for each *test*-panel (i.e. EC reference transcripts vs. *test*-EC panel) are included on each plot. (**B**) The differential correlation score (difference between mean corr. with each set of corresponding reference transcripts) vs. enrichment ranking (position in each respective enriched list, highest correlation = ranking 1) for all transcripts classified as: (i) EC or AC, (ii) EC or OC, (iii) EC or NC, (iv) OC or NC. Threshold lines show ranking below which transcripts were classified as enriched in the annotated cell type.

SUPPLEMENTAL FIGURE 4: **IHC for proteins identified as EC-, AC-, OC- or NC-enriched in human cortex**

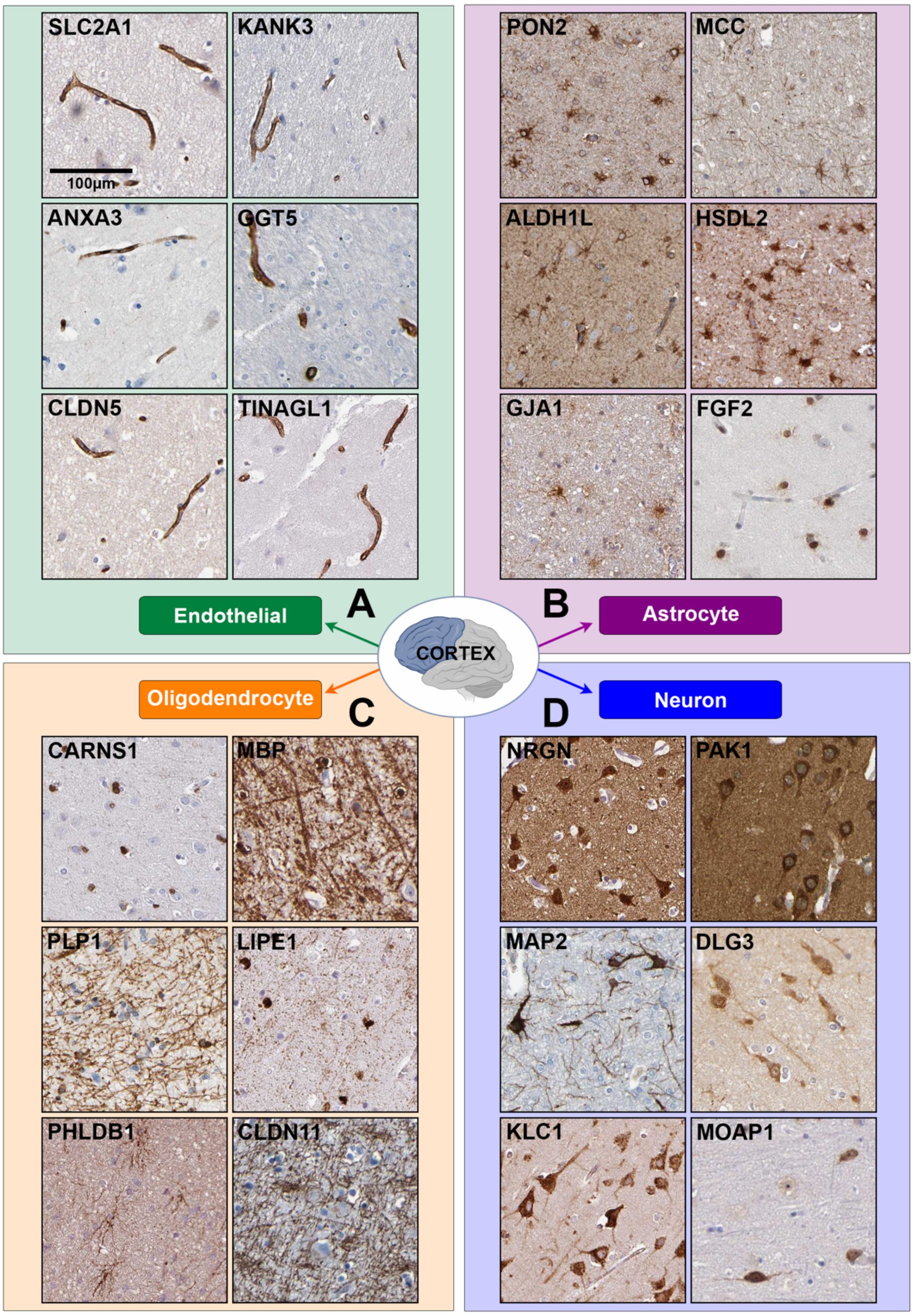


**FIGURE S4. IHC for proteins encoded by transcripts identified as endothelial (EC)-, astrocyte (AC)-, oligodendrocyte (OC)- or neuron (NC)-enriched in human cortex: Related to Figure 3.** Tissue sections from human cortex were stained using primary antibodies targetting proteins encoded by selected transcripts identified as EC-, AC-, OC- or NC-enriched. Scale bar = 100μm

SUPPLEMENTAL FIGURE 5: **Comparison of reference transcript-based and WGCNA for the identification of EC and OC-enriched genes from unfractionated human cortex data**

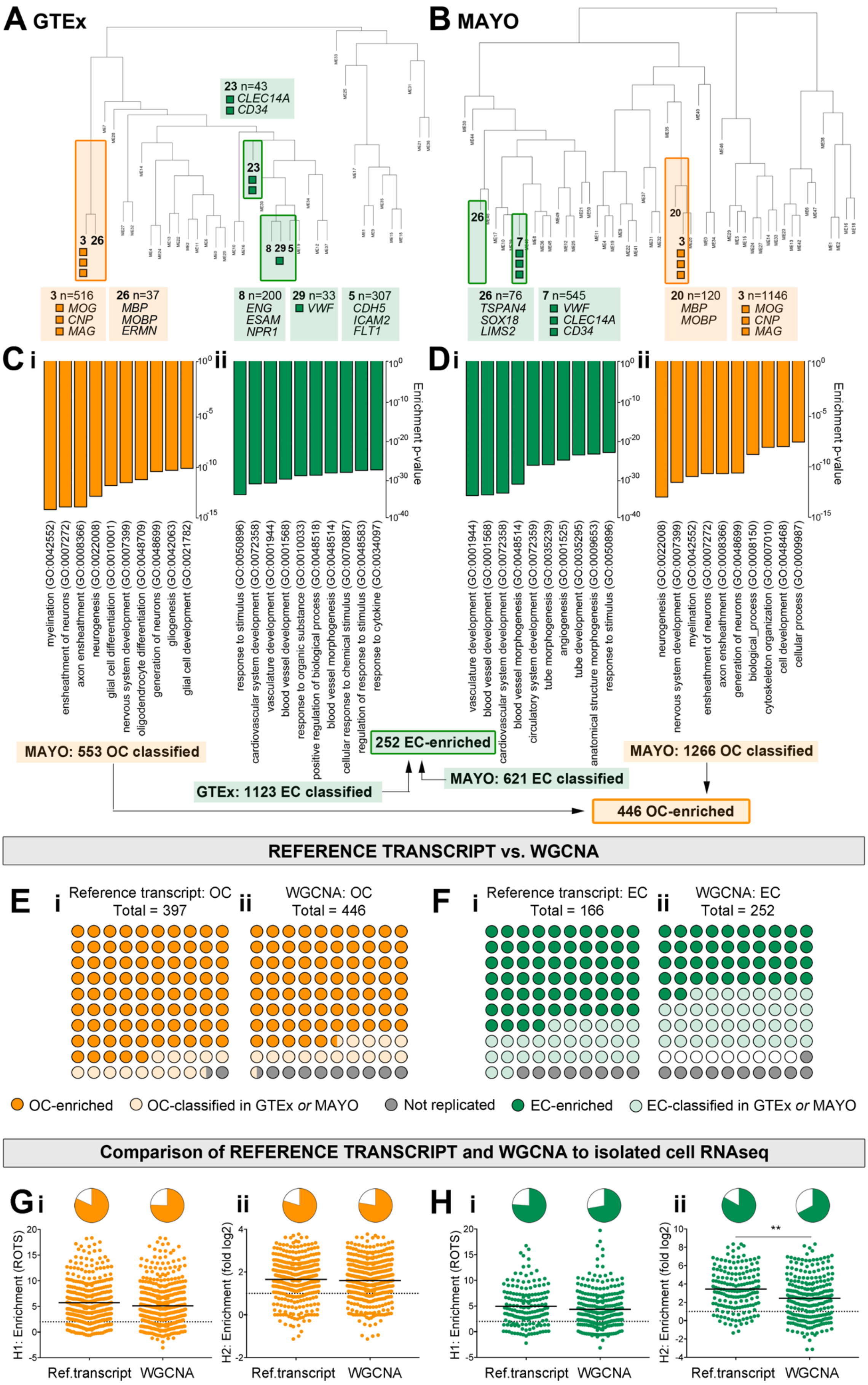

I

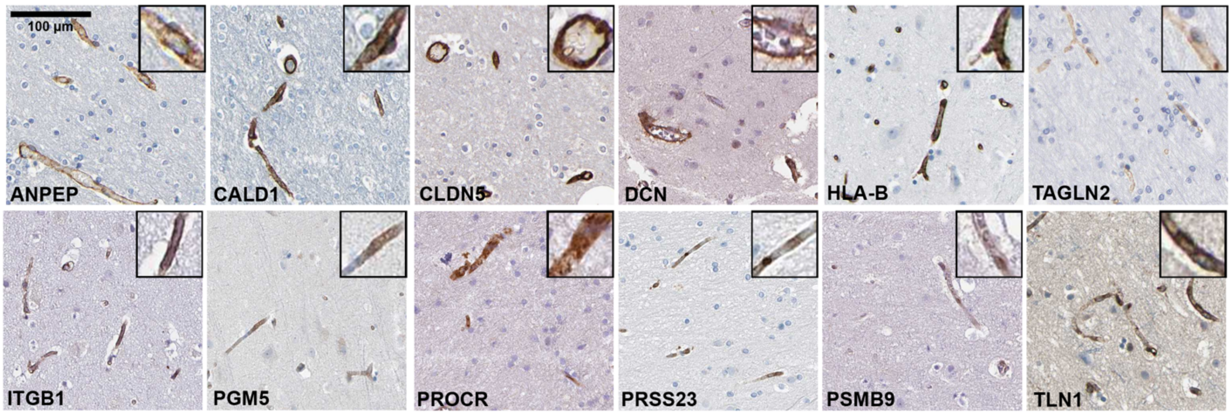


**FIGURE S5. Comparison of reference transcript-based and weighted correlation network analysis for the identification of endothelial- (EC) and oligodendrocyte- (OC) enriched genes from unfractionated human cortex data.** Human cortex RNAseq from the (**A**) Genotype-Tissue Expression [GTEx] portal (n=158) or (**B**) the AMP-AD knowledge portal MAYO RNAseq study (n=80), was subject to a weighted correlation network analysis ('WGCNA') to cluster transcripts into related groups. OC and EC cell-type clusters (highlighted orange and green on dendrogram, respectively) were identified by presence of known corresponding marker transcripts. Enriched gene ontology (GO) biological processes for transcripts classified as (**C**) (**i**) OC- or (**ii**) EC-enriched in GTEx and (**D**) (**i**) EC- or (**ii**) OC-enriched in MAYO. Comparison between analysis methods for transcripts classified as (**E**) OC- or (**F**) EC-enriched in the (**i**) reference transcript vs. WGCNA and (**ii**) WGCNA vs. reference transcript analysis. Datasets from single-cell sequencing of human brain (scRNAseq) (Darmanis et al., 2015) (H1) or RNAseq of isolated cell populations from human brain (H2) (Zhang et al., 2016) were downloaded from the respective publications. Transcript 'enrichment scores' in EC and OC populations in H1 ('ROTS score') and H2 (fold expression vs. other cell types) were compiled for transcripts identified as (**G**) OC- or (**H**) EC-enriched by the (i) reference transcript or (ii) WGCNA analysis. Horizontal line = threshold for enrichment; pie charts - proportion of transcripts reaching threshold. $^{**}p<0.01$ See also Table S6. (**I**) IHC for proteins encoded by transcripts identified as EC-enriched by the reference transcript analysis, but not WGCNA. Tissue sections from human cerebral cortex were stained using primary antibodies targeting proteins encoded by selected transcripts. Scale bar = 100$\mu$m.

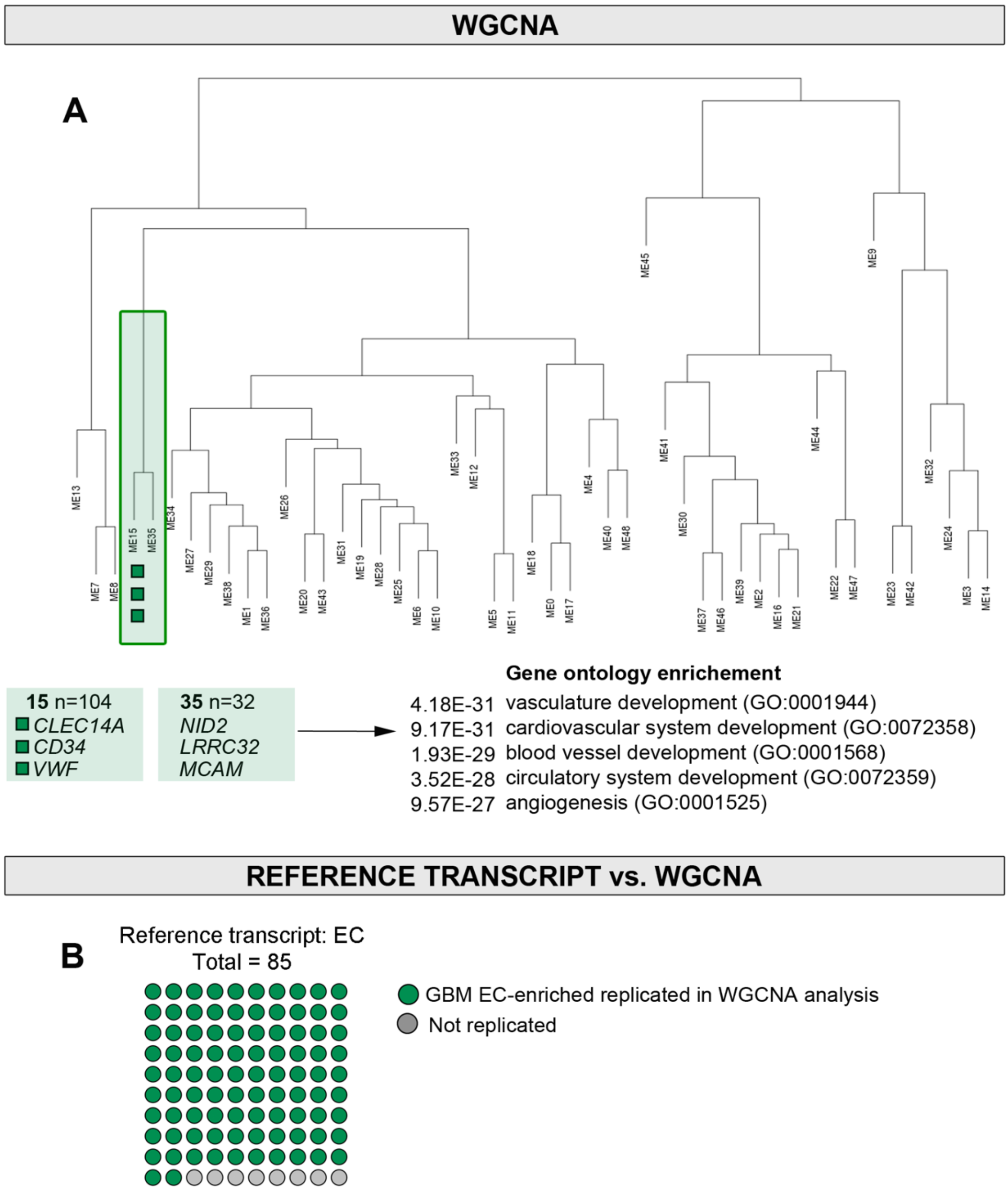


**FIGURE S6. Weighted correlation network analysis identifies endothelial (EC)-enriched genes in unfractionated GBM RNAseq data: Related to Figure 5.** RNAseq data from unfractionated human glioblastoma multiforme (GBM) (n=401) was downloaded from the Cancer Genome Atlas (TCGA) and subject to a weighted correlation network analysis (WGCNA) to cluster transcripts into related groups. (A) EC cell-type clusters (highlighted with green box on dendrogram) were identified by presence of known EC marker transcripts, and the most significantly enriched gene ontology (GO) biological processes for the transcripts in these groups are presented to the right. (B) Replication of transcripts identified as GBM EC-enriched by the reference transcript analysis.

**TABLE S1. Sensitivity and specificity analysis: Correlation coefficient values for test transcript groups in the GTEx (Section A, blue highlighted) and MAYO (Section B, green highlighted) material: Related to Figure 1.** Correlation coefficient values were calculated between the endothelial (EC) reference transcripts *CLEC14A, VWF*, *CD34* (**Tab 1,** 'EC_ref.'), the astrocyte (AC) reference transcripts *BMPR1B*, *AQP4*, *SOX9* (**Tab 2,** 'AC_ref.'), the oligodendrocyte (OC) reference transcripts *MOG*, *CNP*, *MAG* (**Tab 3,** 'OC_ref.'**)** or the neuron reference transcripts *TMEM130*, *STMN2*, *THY1* (**Tab 4,** 'NC_ref.'**)** and the *test*- transcript sets: *Test-EC* (**Table 1**, all tabs), *Test-AC* (**Table 2**, all tabs), *Test-OC* (**Table 3**, all tabs), *Test-NC* (**Table 4**, all tabs), *Test-MG* (**Table 5**, all tabs), and *Test-SMC* (**Table 6**, all tabs). Correlation coefficient values were also calculated between the *test*-transcripts and microglial (MG) reference transcripts *C1QA*, *AIF1* and *LAPTM5* and smooth muscle cell (SMC) reference transcripts *FHL5*, *ACTA2* and *ACTG2* (**Tab 5**, 'MG_SMC_ref.').

**TABLE S2. Full *in vivo* correlation analysis to identify cortex *endothelial (EC)-enriched* transcripts: Related to Figure 1-4.** RNAseq data from unfractionated normal human cortex samples: n=158 from the Genotype-Tissue Expression [GTEx] portal, or n=80 from the AMP-AD knowledge portal (MAYO RNAseq study), were used to generate pair wise Spearman correlation values between **EC** reference transcripts *CLEC14A, VWF* and *CD34* and all mapped protein-coding genes. **Tab 1:** Transcripts fulfilling the EC-enriched analysis criteria (see Figure S2A) (column **A-B**), correlation coefficient values, absolute expression values and false discovery rate (FDR) in GTEx (column **C-K**) or MAYO material (column **L-T**). Classification of gene EC expression from the literature (column **U**), classification of transcript as a core-EC expressed gene in our previous study (Butler et al. 2016) (column **V**). Correlation coefficients for microglial reference transcripts (columns **W-Z**) and SMC reference genes (columns **AA-AD**). **Tab 2:** Full correlation analysis between EC reference transcripts and all mapped protein-coding genes (transcripts with mean FPKM <0.1 in GTEx are excluded), absolute expression values, FDR, Bonferroni corrected p-values and raw p-values in: GTEx (column **C-Q**) or MAYO material (column **R-AF**). **Tab 3:** 100 most highly correlated genes with the EC reference seeds *CLEC14A, VWF* and *CD34* in the MAYO dataset. **Tab 4:** Gene ontology (GO) analysis of transcripts fulfilling EC-enriched analysis criteria: for **Table 1** 'Biological processes': Significant GO terms (column **A**), total number of genes in GO term reference list (entire genome) (column **B**), number of EC-enriched genes in GO term (column **C**), predicted number of

EC-enriched genes in GO term (column **D**), fold enrichment over expected (column **E**), raw p-value (column **F**) and false discovery rate (column **G**). **Table 2** 'Reactome pathways': Significant reactome terms (column **I**), total number of genes in reactome reference list (entire genome) (column **J**), number of EC-enriched genes in reactome pathway (column **K**), predicted number of EC-enriched genes in reactome pathway (column **L**), fold enrichment over expected (column **M**), raw p-value (column **N**) and false discovery rate (column **O**).

**TABLE S3. Full *in vivo* correlation analysis to identify cortex *astrocyte (AC)-enriched* transcripts: Related to Figure 1-4.** RNAseq data from unfractionated normal human cortex samples: n=158 from the Genotype-Tissue Expression [GTEx] portal, or n=80 from the AMP-AD knowledge portal (MAYO RNAseq study), were used to generate pair wise Spearman correlation values between **AC** reference transcripts *BMPR1B, AQP4* and *SOX9* and all mapped protein-coding genes. **Tab 1:** Transcripts fulfilling the AC-enriched analysis criteria (Figure 2SB) (column **A-B**), correlation coefficient values, absolute expression values and false discovery rate (FDR) in GTEx (column **C-K**) or MAYO material (column **L-T**). Correlation coefficients for microglial reference transcripts (columns **U-X**) and SMC reference genes (columns **Y-AB**). **Tab 2:** Full correlation analysis between AC reference transcripts and all other mapped protein-coding genes (transcripts with mean FPKM <0.1 in GTEx are excluded), absolute expression values, FDR, Bonferroni corrected p-values and raw p-values in: GTEx (column **C-Q**) or MAYO material (column **R-AF**). **Tab 3:** 100 most highly correlated genes with the AC reference seeds *BMPR1B, AQP4* and *SOX9* in the MAYO dataset **Tab 4:** Gene ontology (GO) analysis of transcripts fulfilling AC-enriched analysis criteria: for **Table 1** 'Biological processes': Significant GO terms (column **A**), total number of genes in GO term reference list (entire genome) (column **B**), number of AC-enriched genes in GO term (column **C**), predicted number of AC-enriched genes in GO term (column **D**), fold enrichment over expected (column **E**), raw p-value (column **F**) and false discovery rate (column **G**). **Table 2** 'Reactome pathways': Significant reactome terms (column **I**), total number of genes in reactome reference list (entire genome) (column **J**), number of AC-enriched genes in reactome pathway (column **K**), predicted number of AC-enriched genes in reactome pathway (column **L**), fold enrichment over expected (column **M**), raw p-value (column **N**) and false discovery rate (column **O**). **Tab 5**: Transcripts fulfilling EC- and AC-enriched criteria (see Figure 2, Figure S2A and B), with correlation coefficient values, absolute expression values and false discovery rate (FDR) in both GTEx and MAYO for EC and AC reference transcripts.

**TABLE S4. Full *in vivo* correlation analysis to identify cortex *oligodendrocyte (OC)-enriched* transcripts: Related to Figure 1-4.** RNAseq data from unfractionated normal human cortex samples: n=158 from the Genotype-Tissue Expression [GTEx] portal, or n=80 from the AMP-AD knowledge portal (MAYO RNAseq study), were used to generate pair wise Spearman correlation values between **OC** reference transcripts *MOG, CNP* and *MAG* and all mapped protein-coding genes. **Tab 1:** Transcripts fulfilling the OC-enriched analysis criteria (Figure S2C) (column **A-B**), correlation coefficient values, absolute expression values and false discovery rate (FDR) in GTEx (column **C-K**) or MAYO material (column **L-T**). **Tab 2:** Full correlation analysis between OC reference transcripts and all mapped protein-coding genes (excluding transcripts with mean FPKM <0.1 in GTEx), absolute expression values, FDR, Bonferroni corrected p-values and raw p-values in: GTEx (column **C-Q**) or MAYO material (column **R-AF**). Correlation coefficients for microglial reference transcripts (columns **U-X**) and SMC reference genes (columns **Y-AB**). **Tab 3:** 100 most highly correlated genes with the OC reference seeds *MOG, CNP* and *MAG* in the MAYO dataset. **Tab 4:** Gene ontology (GO) analysis of transcripts fulfilling OC-enriched analysis criteria: for **Table 1** 'Biological processes': Significant GO terms (column **A**), total number of genes in GO term reference list (entire genome) (column **B**), number of oligodendrocyte-enriched genes in GO term (column **C**), predicted number of oligodendrocyte-enriched genes in GO term (column **D**), fold enrichment over expected (column **E**), raw p-value (column **F**) and false discovery rate (column **G**). **Table 2** 'Reactome pathways': Significant reactome terms (column **I**), total number of genes in reactome reference list (entire genome) (column **J**), number of OC-enriched genes in reactome pathway (column **K**), predicted number of OC-enriched genes in reactome pathway (column **L**), fold enrichment over expected (column **M**), raw p-value (column **N**) and false discovery rate (column **O**).

**TABLE S5. Full *in vivo* correlation analysis to identify cortex *neuron (NC)-enriched* transcripts: Related to Figure 1-4.** RNAseq data from unfractionated normal human cortex samples: n=158 from the Genotype-Tissue Expression [GTEx] portal, or n=80 from the AMP-AD knowledge portal (MAYO RNAseq study), were used to generate pair wise Spearman correlation values between **NC** reference transcripts *TMEM130, STMN2* and *THY1* and all mapped protein-coding genes. **Tab 1:** Transcripts fulfilling the NC-enriched analysis criteria (Figure S2D) (column **A-B**), correlation coefficient values, absolute expression values and false discovery rate (FDR) in GTEx (column **C-K**) or MAYO material (column **L-T**). Correlation coefficients for microglial reference transcripts (columns **U-X**) and SMC reference genes (columns **Y-AB**). **Tab 2:** Full correlation analysis between NC reference transcripts and all mapped protein-coding genes (transcripts with mean FPKM <0.1 in GTEx are excluded), absolute expression values, FDR, Bonferroni corrected p-values and raw p-values in: GTEx (column **C-Q**) or MAYO material (column **R-AF**). **Tab 3:** 100 most highly correlated genes with the NC reference seeds *TMEM130, STMN2* and *THY1* in the MAYO dataset. **Tab 4:** Gene ontology (GO) analysis of transcripts fulfilling NC-enriched analysis criteria: for **Table 1** 'Biological processes': Significant GO terms (column **A**), total number of genes in GO term reference list (entire genome) (column **B**), number of NC-enriched genes in GO term (column **C**), predicted number of NC-enriched genes in GO term (column **D**), fold enrichment over expected (column **E**), raw p-value (column **F**) and false discovery rate (column **G**). **Table 2** 'Reactome pathways': Significant reactome terms (column **I**), total number of genes in reactome reference list (entire genome) (column **J**), number of NC-enriched genes in reactome pathway (column **K**), predicted number of NC-enriched genes in reactome pathway (column **L**), fold enrichment over expected (column **M**), raw p-value (column **N**) and false discovery rate (column **O**).

**Table S6. Weighted correlation network (WGCNA) analysis identifies endothelial (EC)- and oligodendrocyte (OC)-enriched genes from unfractionated human cortex data: Related to Figure 5.** Human cortex RNAseq from the Genotype-Tissue Expression [GTEx] portal (n=158) or the AMP-AD knowledge portal MAYO RNAseq study (n=80), was subject to a weighted correlation network analysis (WGCNA) to cluster transcripts into related groups. **Tab 1**: Transcripts groupings for GTEx (column **C**) and MAYO (column **D**) datasets. **Tab 2**: Transcript groupings containing the *test*-OC (Table **1**) and *test*-EC (Table **2**) transcripts. **Tab 3**: Lists of: OC-enriched transcripts in GTEx (Table **1**), MAYO (Table **2**) and both (Table **3**) and EC-enriched transcripts in GTEx (Table **4**), MAYO (Table **5**) and both (Table **6**). Transcripts previously identified as core-EC enriched are highlighted green. **Tab 4**: Gene ontology (GO) enrichment in transcript list classified as OC- (Tables **1** and 2) or EC- enriched (Tables **3** and **4**). **Tab 5**: Comparison of the OC- (Table 1 and 2) and EC (Table 3 and 4) enriched transcript lists identified using reference transcript and WGCNA.

**Table S7. Glioblastoma multiforme (GBM) and lower grade glioma (LGG) EC-enriched transcriptomes: Related to Figure 6-8.** RNAseq data from unfractionated human glioblastoma multiforme (GBM) (n=401) and human lower grade glioma (LGG) (n=516) was downloaded from the Cancer Genome Atlas (TCGA) and Spearman's correlation coefficients were calculated between the endothelial (EC) reference transcripts [*CD34*, *CLEC14A*, *VWF*] and all mapped protein-coding genes. **Tab 1:** Full correlation analysis between EC reference transcripts and all other mapped protein-coding genes, correlation coefficient values, mean correlation, and FDR for: GBM (column **C-I**) or LGG (column **J-P**). **Tab 2**: Transcripts identified as EC-enriched in GBM by reference transcript analysis (**Table 1**) or WGCNA (**Table 2**). **Tab 3**: Transcripts identified as EC-enriched in normal cortex, GBM or both. **Tab 4**: Correlation matrices of GBM-specific EC-enriched transcripts in normal (Table **1**) LGG (Table **2**) and GBM (Table **3**)

**Related to FIGURE 1 and TABLE S1:**

19/19 (100%) of *test*-EC genes had correlation values ≥0.52 (mean 0.63 ±0.06, all p<0.001) with the EC reference transcripts [*CD34*, *CLEC14A*, *VWF*] (Figure 1Ai and Table S1, Tab 1: Table 1, Section A). No gene from any other *test*-panel correlated so highly with the EC reference transcripts; 20/20 *test*-AC genes corr. <0.44 (mean corr. 0.26 ±0.12), 20/20 *test*-OC genes corr. <0.13 (mean corr. -0.00 ±0.08), 18/18 *test*-NC genes corr. <0.18 (mean -0.25 ±0.04), 15/15 *test*-MG genes corr. <0.39 (mean 0.25 ±0.10) and 11/11 *test*-SMC genes corr. <0.48 (mean 0.23 ±0.15) (Figure 1Ai and Table S1, Tab 1: Tables 2-6, Section A). 17/20 (90%) of the *test*-AC genes had correlation values ≥0.56 (mean 0.69 ±0.08, all p<0.001) with the AC reference transcripts [*BMPR1B*, *AQP4*, *SOX9*] (Figure 1Aii and Table S1, Tab 1: Table 2, Section A). No gene from any other *test*-panel correlated so highly with the AC reference transcripts; 19/19 *test*-EC genes corr. <0.51 (mean corr. 0.37 ±0.08), all *test*-OC, *test*-NC and *test*-MG category genes corr.<0.31 and all *test*-SMC genes corr. <0.53 (mean corr. 0.11 ±0.20) (Figure 1Aii and Table S1, Tab 2: Tables 1 and 3-7, Section A). 20/20 (100%) of the *test*-OC genes had correlation values ≥0.58 (mean 0.84 ±0.10, all p<0.001) with the OC reference transcripts [*MOG*, *CNP*, *MAG*] (Figure 1Aiii and Table S1, Tab 3: Table 3, Section A). No gene from any other test category correlated so highly with the OC reference transcripts; '*test*-EC, *test*-AC and *test*-MG category genes all corr. <0.27 and all '*test*-NC genes corr. <0.42 (mean corr. 0.14 ±0.15) (Figure 1Aiii and Table S1, Tab 3: Table 3, Section A). 17/18 (94%) of the *test*-NC genes had correlation values ≥0.68 (mean 0.84 ±0.07, all p<0.001) with the NC reference transcripts [*TMEM130, STMN2*, *THY1*] (Figure 1Aiv and Table S1, Tab 4: Table 4, Section A). No gene from any other *test*-panel correlated so highly with the NC reference transcripts; all *test*-EC, *test*-AC, *test*-OC, *test*-MG category genes corr. <0.30 and all *test*-SMC genes corr. <0.50 (mean 0.11 ±0.25) (Figure 1 Aiv and Table S1, Tab 4: Table 1-3, 5-6 Section A).

**Related to FIGURE S5:**

**Identification of OC gene clusters using test-OC transcripts:** The *test*-OC gene panel was used to identify clusters that contained OC markers; in both GTEx and MAYO, 20/20 (100%) of the *test*-OC genes were in 2 clusters (GTEx clusters 3 and 26 and MAYO clusters 3 and 20) (Table S6, Tab 2, Table 1), which were closely related (Figure S5 A and B, orange boxes). In both, the largest cluster contained the OC reference transcripts [*MOG*, *CNP*, *MAG*] and the smaller contained other OC markers e.g. myelin basic protein [MBP], and myelin-associated oligodendrocyte basic protein [MOBP] (Figure S5 A and B). GO enrichment was significant for the biological processes '*myelination*' and '*neurogenesis*' in the GTEx and MAYO clusters (Figure 5Ci and Dii) (Table S6, Tab 4, Table 1-2). 446 transcripts were classified as OC-enriched in GTEx *and* MAYO data, constituting the final list (Table S6, Tab 3, Column I-L).

**Identification of EC gene clusters using test-EC transcripts:** The *test*-EC gene panel was used to identify clusters containing EC markers; in the GTEx analysis, 18/19 (95%) of the *test*-EC genes were distributed between 4 clusters (GTEx 5, 8, 23 and 29) (Figure 5A, green boxes) (Table S6, Tab 2, Table 2). EC reference transcripts [*CD34*, *CLEC14A*, *VWF*] were divided between 2 clusters (Figure S5 A). All 4 contained transcripts we previously identified as body-wide core EC-enriched (Butler et al., 2016); cluster 5: 21/307 (7%), cluster 8: 40/200 (20%), cluster 23: 16/43 (37%) and cluster 29 9/33 (27%) (Table S6, Tab 3, Column N-P, highlighted green). In the MAYO data, 16/19 (89.5%) of the *test*-EC were in a single group (cluster 7), which contained the EC reference transcripts [*CD34*, *CLEC14A*, *VWF*], and 2/19 (10.5%) were in an additional smaller group (cluster 26) (Figure 5B) (Table S6, Tab 2, Table 2). Both groups contained transcripts previously identified as body-wide core EC-enriched (Butler et al., 2016); cluster 7: 89/545 (16%), cluster 26: 16/76 (20%) (Table S6, Tab 3, Column R-T, highlighted green). GO enrichment was significant for vascular-related biological processes, including '*cardiovascular system development*' and '*blood vessel morphogenesis*' (Figure 5Cii and Di) (Table S6, Tab 4, Table 3-4). 252 transcripts were classified as EC-enriched in GTEx *and* MAYO data, constituting the final list (Table S6, Tab 5, Column V-Y).

**Reproducibility of results between reference transcript analysis and WGCNA methods:** 337/397 [85%] of genes classified as OC-enriched by the reference transcript analysis were also identified by WGCNA (Figure S5 Ei). Of those not replicated, 55/60 (92%) were classified as OC-enriched in either the GTEx *or* MAYO WGCNA, but not *both*, and thus were excluded (Figure S5 Ei) (Table S6, Tab 5, Columns A-D). Correspondingly, 337/446 [76%] of genes classified as OC-enriched by the WGCNA were also identified by the reference transcript analysis (Figure S5 Eii). Of those not replicated, 67/109 (61%) were classified as OC-enriched in either the GTEx *or* MAYO reference transcript analysis, but not in *both* (Figure 5Eii) (Table S6, Tab 5, Columns F-I). In most cases, values narrowly failed to meet the required threshold in the 'negative' dataset (mean corr. GTEx 0.57 ±0.07, MAYO 0.75 ±0.08) (threshold required: ≥0.60 in GTEx, ≥0.70 in MAYO). The remaining 42/109 (39%) of those not replicated failed to reach the threshold for OC-enriched classification in either the GTEx or the MAYO reference transcript analyses, but by a narrow margin (mean corr. GTEx 0.53 ±0.06, MAYO 0.62 ±0.07). 106/166 [64%] of genes classified as EC-enriched by the reference transcript analysis were also identified by the WGCNA (Figure S5 Fi). Of those not replicated, 47/60 were classified as EC-enriched in the either the GTEx *or* MAYO WGCNA, but not in *both*, and thus were excluded (Figure 5Fi) (Table S6, Tab 5, Columns L-O). GO enriched terms in this list included '*circulatory system development*' (p<7.98E-08), '*blood vessel morphogenesis*' (p<4.8E-7) and '*angiogenesis*' (p<1.0E-5), and included known EC-genes (Butler et al., 2016), many of which we confirmed as EC-enriched by IHC (Figure S5). Thus, these are likely false negatives in the WGCNA. The WGCNA identified more EC-enriched genes than the reference transcript (252 vs. 166), and 106/252 (42%) of genes classified as EC-enriched by the WGCNA were also identified by the reference transcript analysis (Figure S5 Fii) (Table S6, Tab 5, Columns Q-T). Of those not replicated, 23/146 (16%) reached the required correlation threshold for classification as EC-enriched in the reference transcript analysis, but were excluded due to high correlation values with microglia-enriched (MG) transcripts (n=9) or dual-enrichment with AC (n=14), while 29/146 (20%) had correlations with EC reference transcripts <0.05 below the designated threshold for classification (mean corr. GTEx 0.49 ±0.04, min. 0.45, mean corr. MAYO ≥0.59 ±0.08, min 0.46).
